# A molecular phylogeny of forktail damselflies (genus *Ischnura*) reveals a dynamic macroevolutionary history of female colour polymorphisms

**DOI:** 10.1101/2020.06.06.137828

**Authors:** Rachel Blow, Beatriz Willink, Erik I. Svensson

## Abstract

Colour polymorphisms are popular study systems among biologists interested in evolutionary dynamics, genomics, sexual selection and sexual conflict. In many damselfly groups, such as in the globally distributed genus *Ischnura* (forktails), sex-limited female colour polymorphisms occur in multiple species. Female-polymorphic species contain two or three female morphs, one of which phenotypically matches the male (androchrome or male mimic) and the other(s) which are phenotypically distinct from the male (heterochrome). These female colour polymorphisms are thought to be maintained by frequency-dependent sexual conflict, but their macroevolutionary histories are unknown, due to the lack of a robust molecular phylogeny. Here, we present the first time-calibrated phylogeny of *Ischnura*, using a multispecies coalescent approach (StarBEAST2) and incorporating both molecular and fossil data for 41 extant species (55% of the genus). We estimate the age of *Ischnura* to be between 13.8 and 23.4 millions of years, i.e. Miocene. We infer the ancestral state of this genus as female monomorphism with heterochrome females, with multiple gains and losses of female polymorphisms, evidence of trans-species female polymorphisms and a significant positive relationship between female polymorphism incidence and current geographic range size. Our study provides a robust phylogenetic framework for future research on the dynamic macroevolutionary history of this clade with its extraordinary diversity of sex-limited female polymorphisms.

## Introduction

Colour polymorphisms are widespread among many animal taxa and provide unique opportunities to study the evolutionary processes shaping genetic diversity within- and between-populations. The genetic basis of such colour polymorphisms is generally due to multiple segregating alleles at a single locus, with different degrees of dominance in relation to each other (Svensson, 2017; Svensson et al., 2009; Wellenreuther et al., 2014). Population-level studies aim to disentangle the evolutionary mechanisms responsible for the coexistence of alternative morphs in sympatry (Le Rouzic et al., 2015; Svensson et al., 2005). Yet, the turnover of alternative phenotypes and their underlying alleles on a macroevolutionary scale is more poorly understood (Corl et al., 2010; Jamie and Meier, 2020). Phylogenetic comparative methods can be used to explore how discrete phenotypic polymorphisms arise and are maintained or lost across macroevolutionary time scales (Corl et al., 2010; Hugall and Stuart-Fox, 2012; Jamie and Meier, 2020) . However, these methods are reliant on robust phylogenetic inferences (Rangel et al., 2015), and are affected by the incidence of evolutionary transitions between character states (Uyeda et al., 2018). An important step to elucidate how microevolutionary processes influence the macroevolutionary fate of discrete and alternative phenotypes is therefore to reconstruct, expand and improve phylogenies of clades with a dynamic history of phenotypic polymorphisms.

Discrete and heritable polymorphisms such as colour morphs are often sex-limited in their expression (Neff and Svensson, 2013; Svensson et al., 2009). Whereas male colour polymorphisms are typically associated with different strategies used in competition over access to females (Neff and Svensson, 2013), female-limited colour polymorphisms have been linked either to predator-avoidance through interspecific mimicry (Kunte, 2009), avoidance of male mating harassment (Gosden and Svensson, 2009; Lee et al., 2019; Neff and Svensson, 2013) or a combination of these two factors (Cook et al., 1994). In the insect order Odonata (dragonflies and damselflies), female-limited colour polymorphisms occur in several genera and many species (Fincke et al., 2005). Female colour polymorphisms are particularly common in the globally distributed genus of pond damselflies *Ischnura* (family Coenagrionidae; Fig. 1) (Gering, 2017; Gosden and Svensson, 2009; Sirot and Brockmann, 2001; Takahashi and Watanabe, 2010). Within *Ischnura,* there are both female-monomorphic (FM) and female-polymorphic (FP) species, but males are always monomorphic (Fincke et al., 2005; Sanchez-Guillen et al., 2011; Willink et al., 2019)(Fig. 1B). Of the approximately 75 species in the genus *Ischnura*, c.a. 40% are FM, 40% are FP, and the female-colour status of the remaining 20% is currently unknown (Willink et al., 2019).

**Figure 1.**
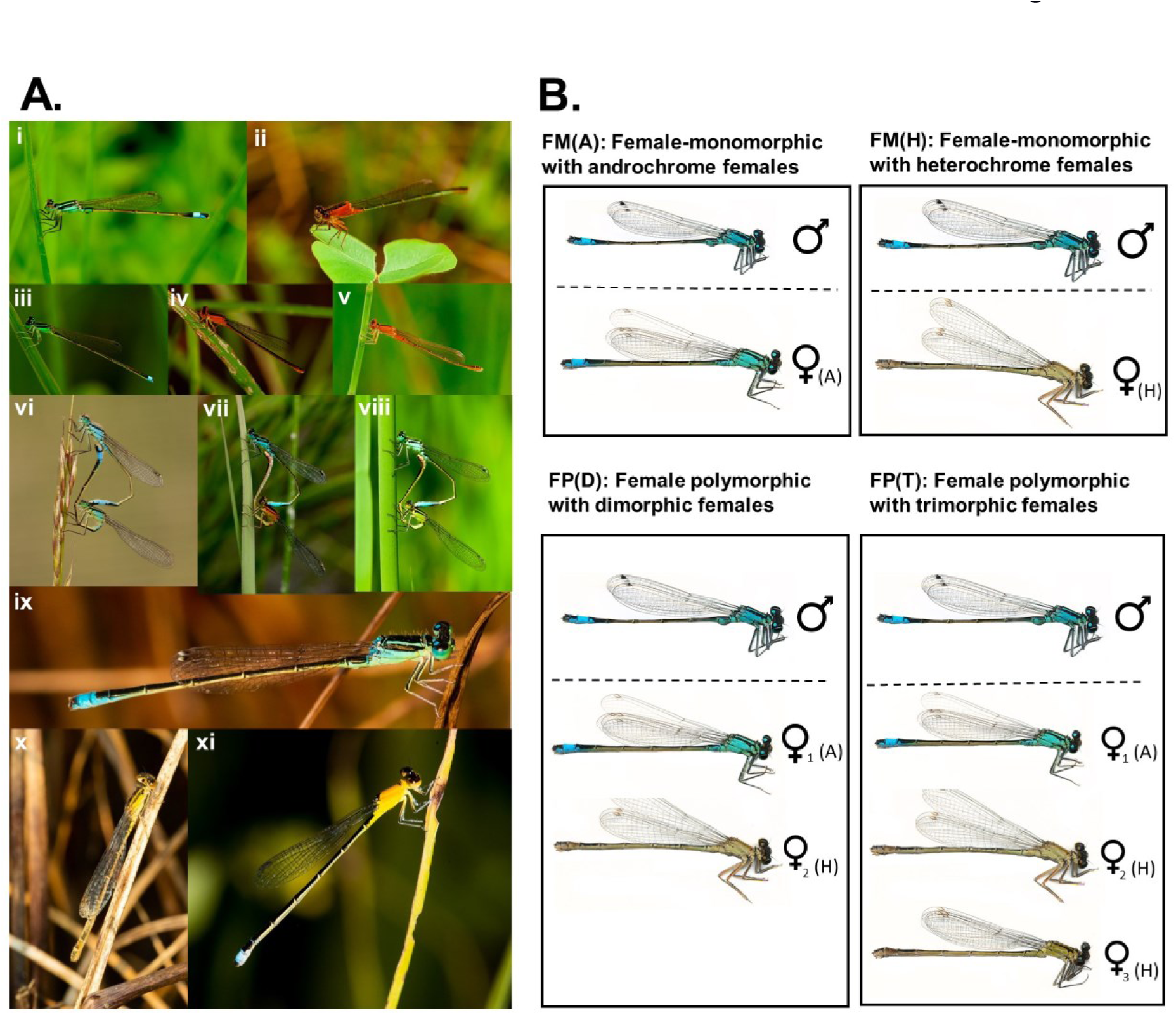
Phenotypic variation in the damselfly genus *Ischnura.* **A.** Some representative species, from both the Old and the New World, exemplifying the extensive phenotypic and colour variation within and between species in the genus *Ischnura,* as well as age-related variation (note that the orange and yellow females in **ii**, **iv**, **v** and **xi** show sexually immature and age-related colour phases expressed in females only). All species in this panel are included in the molecular phylogeny we present in this paper. **i.** Sexually mature male and **ii.** immature female *I. ramburii* (Texas, USA). **iii.** Sexually mature male and **iv.** immature female *I. senegalensis* (Japan). **v.** immature female *I. pumilio* (Sweden). **vi-viii:** Some species of *Ischnura* also show heritable female colour polymorphism at the adult stage. Here are the three different female colour morphs in the trimorphic species *I. elegan*s (all photos from Sweden). **vi.** Male *I. elegans* mating with an androchrome (male mimic) female morph. **vii.** Male *I. elegans* mating with a heterochrome morph (*infuscans-obsoleta*) . **viii.** Male *I. elegans* mating with the other heterochrome morph (*infuscans*). The sister species of *I. elegans* is *I. graellsii* (photos from Portugal): **ix.** sexually mature male and **x.** sexually mature heterochrome female morph (*infuscans*). Finally, **xi** shows an immature female *Ischnura capreolus*, revealing the yellow colour females of this species exhibit prior to sexual maturation (Brazil). All photographs taken by E.I. Svensson. **B.** The four possible female colour morph states found in the damselfly genus *Ischnura*. Here, we illustrate these four colour morph states using scans taken from *I. elegans.* Female-monomorphic (FM) species do not exhibit any discrete intrasexual female colour variation. However, in all polymorphic *Ischnura* species, emales are either phenotypically similar to males (androchrome) (FM(A)) or phenotypically distinct from males (heterochrome) (FM(H)). Female-polymorphic (FP) species have multiple female colour morphs. Polymorphic *Ischnura* species are either dimorphic (FP(D): one androchrome and one heterochrome female morph, two morphs in total) or they are trimorphic (FP(T): one androchrome and two heterochrome female morphs, three morphs in total).

In FP species, females are either similar in colouration (Fig. 1B) and sometimes also in body shape to males, here referred to as androchrome females or male mimics, or they are phenotypically different from males (referred to here as heterochrome females) (Gosden and Svensson, 2009). Amongst these FP taxa, some species are female-dimorphic (FP(D)) with one androchrome and one heterochrome female morph, whereas others are female-trimorphic (FP(T)), with one androchrome and two distinct heterochrome female morphs (Fig. 1B). In FM species of *Ischnura*, both males and females are therefore monomorphic (Willink et al., 2019)(Fig. 1B). A species can therefore be either sexually monomorphic with androchrome females (FM(A)) or sexually dimorphic with heterochrome females (FM(H))(Willink et al., 2019)(Fig. 1B). Such interspecific variation in female colour states is particularly useful for phylogenetic comparative studies on the macroevolutionary dynamics of discrete and heritable polymorphisms (Corl et al., 2010; Jamie and Meier, 2020).

The classical explanation for the evolutionary origin and adaptive function of female colour polymorphisms in *Ischnura,* and other damselflies is the “male-mimicry hypothesis” (Cordero, 1989; Robertson, 1985). Male-mimics are thought to have evolved as a response to selection from sexual conflict over mating, which favoured visual features that made such male-like females less recognizable as potential mates to males (Cordero et al., 1998; Gosden and Svensson, 2009; Willink et al., 2019; Xu et al., 2014). This mimicry hypothesis would phylogenetically predict that the common ancestor of FP damselflies was an FM(H) species, in which a novel male-mimicking female phenotype emerged and subsequently increased in frequency due to a frequency-dependent fitness advantage of male mimicry. An alternative, though not mutually exclusive hypothesis for the evolution and maintenance of such female polymorphisms are flexible search images in males (Fincke, 2004; Miller and Fincke, 1999). Under this alternative hypothesis, males are expected to switch mating preferences between female morphs, depending on their frequency, resulting in increased male preferences for currently common morphs (Van Gossum et al., 2001). The most common female morph is then expected to be disproportionally harassed, resulting in negative frequency-dependent selection and the stable maintenance of female polymorphism across many generations (Gosden and Svensson, 2009; Iserbyt et al., 2013, 2011; Le Rouzic et al., 2015; Svensson et al., 2005; Takahashi et al., 2014, 2010). Unlike a pure version of the male-mimicry hypothesis (described above), female colour polymorphisms in *Ischnura* could under this latter scenario have evolved from either an FM(H) or FM(A) ancestral state (Fig. 1B).

FM species of *Ischnura* have received less attention compared to FP taxa, and consequently we know very little about why there is such a striking variation in the presence or absence of female polymorphisms in different subclades within this genus. One explanation could be that FM lineages reflect an ancestral state with low intensity of sexual conflict. If so, female polymorphisms may have subsequently evolved in lineages experiencing novel selective regimes of more intense sexual conflict. This could occur, for instance, due to a rapid increase in population density or transition to a new mating system. Alternatively, FM lineages could reflect transitions from an ancestral state of female polymorphism (FP), via founder effects or genetic drift, or through the selective fixation of one of the alternative previously sympatric morphs (Corl et al., 2010; West-Eberhard, 1986). Such speciation by morph loss has been termed “morphic speciation” (Corl et al., 2010; West-Eberhard, 1986). The morphic speciation hypothesis explicitly predicts that monomorphic taxa should primarily be located at the terminal tips on a phylogeny, with polymorphism being the ancestral state (Corl et al., 2010). Regardless of the generality and validity of the morphic speciation hypothesis, our knowledge about the relationship between polymorphism and speciation is limited and there are only a few phylogenetic comparative studies in this area (Hugall and Stuart-Fox, 2012; McLean and Stuart-Fox, 2014; Schwander and Leimar, 2011). Recently, it has also been emphasized that some colour polymorphisms can be maintained across speciation events resulting in trans-species polymorphisms (Jamie and Meier, 2020).

There have been a few previous comparative studies in the genus *Ischnura,* aiming to understand the ecological correlates of female colour polymorphisms and their evolutionary origin. These previous studies suggested that female colour polymorphism was the ancestral state in *Ischnura* (Mattern and Van Gossum, 2008; Sánchez-Guillén et al., 2020). They further argued female polymorphisms are more common in species with female polyandry, than in species with (putative) female monandry, consistent with female polymorphism being promoted by sexual conflict (Mattern and Van Gossum, 2008; Robinson and Allgeyer, 1996; Sánchez-Guillén et al., 2020). However, these previous comparative studies suffer from several limitations, such as ignoring the shared evolutionary history of closely related species

(Robinson and Allgeyer, 1996), using highly sensitive methods of ancestral state reconstruction like maximum parsimony(Mattern and Van Gossum, 2008), and ignoring phylogenetic and model uncertainty in ancestral state estimation (Sánchez-Guillén et al., 2020). Ignoring both phylogenetic uncertainty, model uncertainty and using highly sensitive methods like maximum parsimony is problematic, particularly for rapidly evolving characters (Cunningham et al., 1998; Schluter et al., 1997), which is likely the case for these colour polymorphisms and other traits that are involved in sexual conflict (Rice and Holland, 1997). Here, we revisit these previous hypotheses for the evolution of female colour polymorphism in *Ischnura*, with a broadened taxon sampling in our phylogenetic inference, and using contemporary methods to simulate trait evolution under a Bayesian framework. This approach allows for model-based inference of ancestral states, while taking into account model and phylogenetic uncertainty (Pagel et al., 2004; Pagel and Meade, 2006).

Our novel phylogenetic and comparative data and analyses in this study aimed to shed light on the origin of female colour polymorphisms. We also estimate the age and study the evolutionary history of the genus *Ischnura*. We use a multispecies coalescent model to simultaneously estimate a posterior distribution of species tree topologies and divergence times for 55% of species in the genus *Ischnura.* We then investigate the macroevolutionary history of female colour polymorphisms by reconstructing the ancestral state of this trait. Finally, we investigated whether the presence of female polymorphisms are associated with large geographic range sizes, a factor that is likely to reflect population size and the potential for morphic speciation by neutral processes. A positive association between the presence of female colour polymorphisms and geographic range would be expected in both of the following scenarios: 1) if morphs are lost by neutral or selective processes after a reduction of population (and range) size or 2) if the effects of female colour polymorphisms on population fitness allow FP species to sustain larger populations over a greater geographic range.

Our study contributes to increasing our understanding of the macroevolutionary history of this genus, the origin of female colour polymorphism and the demographic factors promoting the maintenance of these female polymorphisms. In presenting these results, we hope to stimulate future empirical investigations on these and other damselfly clades, and we discuss some promising hypotheses for follow-up studies. We note that many damselfly groups, although popular in much ecological and evolutionary research (Cordoba-Aguilar, 2008) are still relatively poorly investigated with respect to phylogenetic relationships and evolutionary history, although some recent progress has been made, particularly within the family Coenagrionidae (Beatty et al., 2017; Callahan and McPeek, 2016; Dijkstra et al., 2014; Swaegers et al., 2014; Torres-Cambas et al., 2019; Vega-Sánchez et al., 2019).

## 2. Materials and Methods

### 2.1 Data collection

We have gathered DNA-sequence data, phenotypic data and geographic range size data for 41 species of *Ischnura*. Two of the taxa in this dataset are nominally in the genus *Pacificagrion*, however recent molecular data shows they are nested within the *Ischnura* clade (Willink et al. 2019). Likewise, two other taxa, including a yet undescribed species, were classified in the genus *Amorphostigma* (Marinov et al., 2015). However, Dijkstra et al. (2014) subsumed *Amorphostigma* into *Ischnura*, and recent molecular data support this change (Willink et al. 2019). Here, we use the generic names *Ischnura* and *Pacificagrion*, according to the “World Odonata List” (https://www.pugetsound.edu/academics/academic-resources/slater-museum/biodiversity-resources/dragonflies/world-odonata-list2/), which maintains a taxonomic classification of Odonata with currently valid names.

We obtained sequences for three nuclear and two mitochondrial markers. The nuclear markers include conserved and hypervariable regions of the 28S ribosomal DNA (D7; alignment length: 693 bp), coding and intronic regions of the arginine methyltransferase gene (PMRT; alignment length: 544 bp), and a coding region of the histone 3 (H3; fragment length: 328 bp). The mitochondrial markers sampled were the third domain of the mitochondrial 16S ribosomal DNA (16S; alignment length: 397 bp) and a fragment of the cytochrome oxidase subunit I (COI; fragment length: 481 bp). Sequence data was collected for 147 specimens, of which 64 were sequenced for this study (Table S1). Genomic DNA was extracted from 1-2 legs using a Qiagen DNeasy tissue kit and following the manufacturer’s instructions, except that we increased the amount of proteinase K and incubation time for museum samples collected before 2005, in order to obtain a higher DNA yield. Polymerase chain reaction (PCR) primers and amplification programs used for the molecular markers were the same as in Willink et al. (2019). Amplification products were purified using the Affymetrix ExoSAP-IT reagent (Cat. No. 78200) and sequenced on an ABI 3100 capillary sequencer at Lund University. These novel sequence data were complemented with already-published sequences (Table S1), downloaded from the National Centre for Biotechnology Information (https://www.ncbi.nlm.nih.gov/genbank/). There were no indels in the sequences of two molecular markers (COI and H3), which were aligned using MUSCLE v. 3.8.31 under default settings (Edgar, 2004). To reduce gap overmatching in ribosomal sequences (16S and 28S) and intronic regions (present in PMRT), we used the phylogenetically informed algorithm PRANK v.150803 (Löytynoja and Goldman, 2008, 2005) to align these data. PRANK was iterated 20 times, inferring a new guide tree after each alignment and the highest scoring alignment was selected by randomly breaking score ties at each iteration. The remaining settings were left in their default values, except for the alignment of 28S rDNA, for which a branch length multiplier of two was used to produce more mismatches (fewer gaps) in the hypervariable region.

Data on female colour morph status (i.e. FM, or FP) has been recently compiled for *Ischnura* damselflies, based on published literature, field observations, examination of museum specimens and online resources (Willink et al., 2019). We complemented these data with information on whether FP species had two or three morphs (i.e. FP(D) or FP(T)) and whether FM species had androchrome or heterochrome females (i.e. FM(A) or FM(H)) (Table S2). Here, species with a single female morph were classified as FM(A), if they had only one female morph with colouration and patterning similar to that of conspecific males, or as FM(H), if they had only one female morph with colouration and patterning markedly different from the conspecific males. If a species contained two distinct female morphs (always one androchrome and one heterochrome), we classified it as FP(D). Finally, species that had three morphs (which were always one androchrome and two heterochrome female morphs), were classified as FP(T).

We complemented these colour-state data with coarse-grained estimates of geographic range size (Table S2). Because accurate geographic distribution data is lacking for many species, we estimated geographic range using occurrence data from administrative regions, mainly countries, and where available, states or provinces (Table S2). Presence data was translated to geographic range size (km^2^) by adding the areas of all the countries or other administrative regions where a species is known to occur (Table S2), as has been done previously in comparative studies where geographic range is taken as a proxy for population size (Ross et al., 2013). The areas of administrative regions were extracted from the GADM database of global administrative areas v. 3.6 (Global Administrative Areas, 2018).

### 1.2 Time-calibrated species tree

We analysed sequence data under a multispecies coalescent (MSC) model using StarBEAST2 (Ogilvie et al., 2017). The MSC models the evolution of gene trees independent from each other but conditional on a shared species tree (Degnan and Rosenberg, 2009). When implemented in a Bayesian framework, the MSC provides a means to estimate the joint conditional distribution of gene trees and species trees in the face of gene tree discordance due to incomplete lineage sorting (Rannala et al., 2020). Incomplete lineage sorting occurs whenever two alleles in the same population fail to coalesce, going backwards in time, before at least one of them coalesces with an allele from a more distantly related population. Such incomplete lineage sorting is commonplace, particularly during rapid radiations, such as the one described in the closely related damselfly genus *Enallagma* (Callahan and McPeek, 2016). By using the MSC model, we allowed genealogical histories to vary among nuclear loci and between nuclear and mitochondrial genes. As there is no recombination amongst mitochondrial genes, we linked the gene tree models of mitochondrial sequences (COI and 16S).

We placed a birth-death prior, conditional on the root, on the species tree to allow for extinction, set the gene ploidy to 0.5 for mitochondrial sequences and 2.0 for nuclear sequences and allowed population-size to be analytically integrated to avoid over-parameterization. We used a relaxed uncorrelated log normal (UCLN) clock per locus to allow for variation in evolutionary rates between lineages. Substitution models for each locus were inferred during the analysis using the package “bModelTest” (Bouckaert and Drummond, 2017) to allow for site model uncertainty. We placed a lognormal prior with a mean of 0.0115 and standard deviation of 1.0 on each relaxed mitochondrial clock and a broader lognormal prior with a mean of 0.0115 and standard deviation of 2.0 on each relaxed nuclear clock to centre clock rates on the suggested insect mitochondrial divergence rate of 2.3% per million years (Brower, 1994) whilst allowing for clock rate uncertainty, particularly in the nuclear loci.

### Data for *Ischnura* fossils were downloaded from the Paleobiology database (www.pbdb.org)

on 14^th^ May 2018. There were three fossils classified as either in or closely related to the genus *Ischnura*. Two of these fossils were relatively young (7.2 - 5.3 mya), each one consisting of a single wing with missing fragments (Gentilini and Bagli, 2004). The third fossil was preserved in Dominican amber (20.44 – 13.82 mya) and was identified and described as *Ischnura velteni* based on a combination of wing venation and morphological traits in other body parts (Bechly, 2000). The combination of these characters suggests that *I. velteni* belongs to the extant genus *Ischnura* (Bechly, 2000), and we therefore considered it the most appropriate crown fossil with which to calibrate the root age of the *Ischnura* tree. We placed an exponentially distributed prior on the root age of the species tree with an offset of 13.8 to ensure the most recent common ancestor (MRCA) of *Ischnura* could not have appeared after the minimum age of this known member of the genus. The mean of the exponential was drawn from a gamma hyperprior with a shape parameter α = 5.0 and an inverse scale parameter β = 2.0. This hyperprior specification centred the root age prior distribution around 10 mya prior to the offset, allowing a moderately high level of uncertainty in the age difference between the fossil and the MRCA of *Ischnura*.

We ran two separate Markov Chain Monte Carlo (MCMC) algorithms for 200 million iterations with a sampling rate of 20000. We confirmed that independent chains were stationary and had converged for these and all subsequent analyses in R v.3.6.1 (R Core Team 2019) using the “coda” package (Plummer et al., 2006). We combined posterior trees using LogCombiner (available as part of BEAST v2.4.7)(Bouckaert et al., 2014), simultaneously discarding 20% of the posterior as burnin, and summarized them into a maximum clade credibility (MCC) tree using TreeAnnotator (available as part of BEAST v2.4.7). The MCC tree and all further phylogenetic plots were annotated in R v.3.6.1 (R Development Core Team, 2019) using the packages “ape” (Paradis and Schliep, 2019), “ggtree” (Yu et al., 2017) and “ggplot2” (Wickham, 2016).

### 1.3 Reconstruction of ancestral character states of female colour polymorphism

We used the Multistate-method in BayesTraits v.3.0 (Pagel et al., 2004) to infer female-colour character states at ancestral nodes, given this new *Ischnura* phylogeny and the female-colour character states of extant species. To account for phylogenetic uncertainty, we ran the model using a sample of 1000 trees from the combined species-tree posterior of the two independent runs. Additionally, we ran the model using reversible jump (RJ) MCMC to account for uncertainty in the number of model parameters. In this model, the number of different transition rate parameters and their values are sampled in proportion to their posterior probability. Consequently, in each posterior sample a number of transition rates are set to zero, and the non-zero rates belong to one or more categories. We placed an exponential prior on all rate parameters, with the mean seeded from a uniform hyperprior bounded between 1 and 100. We ran two separate chains for 10 million iterations, sampling every 1000 with 20% posterior burnin. Trees were scaled to an average branch length of 0.01 to prevent various rates from becoming too small. Although our main focus is on ancestral state reconstructions, we present the posterior distributions of transition rate parameters in the Supporting Material (Fig. S1).

A recent study estimated ancestral female-colour states, using a single MCC tree and stochastic character mapping (Sánchez-Guillén et al., 2020). We investigated if the conflicting results between the present study and Sánchez-Guillén et al. (2020) could be explained by differences in taxon sampling. Specifically, we examined if inclusion of taxa sampled by Sánchez-Guillén et al. (2020), but not included in the present study would qualitatively change our conclusions.

To do so, we used the R package “pastis” (Thomas et al., 2013) to include the missing taxa in our MCC species-tree, by randomly assigning their phylogenetic placement and branch lengths within their inferred clade, according to Sánchez-Guillen et al. (2020) (See Supporting Methods). We then repeated ancestral state inferences as explained above on the sample of expanded species trees.

### 1.4 Linking female colour polymorphism to geographic range size

We investigated if the occurrence of female colour polymorphisms could be influenced by demographic changes, by using geographic range size of extant species as a proxy for population size. Geographic range size is also a biologically interesting variable in the light of recent work showing that female colour polymorphisms can increase population fitness (Takahashi et al., 2014). We used a Bayesian phylogenetic mixed-effect model (BPMM) (Hadfield and Nakagawa, 2010) implemented in the package MCMCglmm v. 2.29 (Hadfield, 2010) to test for such phylogenetic correlation. Female-morph status was treated as a binary response variable, with species being classified as either FM or FP. Since the occurrences of FP(T) species and FP(A) were low, we classified these species-states as being FP and FM, respectively. In this analysis, we thus investigated if the probability of a species being FP was significantly and positively related to geographic range size (a continuous predictor variable).

We accounted for phylogenetic uncertainty by using a random sample of 2100 species-trees generated in StarBEAST2. To estimate the phylogenetic random effect, each tree was sampled for 10000 iterations, of which only the last iteration was saved to the posterior, and the first 100 trees were also discarded as burn in (O’Connor et al., 2018). We used a Knonecker prior (mu=0, V= σ2units + π^2/3^) for the fixed effects as it is relatively flat when a logit link function is used (Hadfield, 2015). We used a parameter-expanded χ^2^ distribution (V=1 nu=1000, alpha.mu=0, alpha.V = 1) for the prior on the phylogenetic variance components (Villemereuil et al., 2013). The residual variance was fixed to 1, as it cannot be identified for binary response models. As a statistical test of whether the probability of female colour polymorphism increases with range size, we report the fraction of the posterior distribution in which increasing range size does not result in an increasing probability of female colour polymorphism. The results of this analysis were visualized by plotting the expected probability of female colour polymorphism against estimated geographic range size.

## 2. Results

### 2.1 Divergence time estimation

Our time-calibrated phylogeny suggests that the age of the MRCA of the genus *Ischnura* lies in a 95% highest posterior density (HPD) interval between 13.8 and 23.4 mya, i.e. during the early Miocene (Fig. 2). A comparison of the posterior root age in our model to a model ran without molecular data suggests the sequence data were sufficiently informative to narrow this posterior distribution (Fig. 3), although age estimates had quite broad error margins (Fig. 2). Nonetheless, divergence dating revealed several interesting features, such as the very recent divergence of two of the most intensely studied FP species in Europe (*I. elegans* and *I. graellsii*) (Fig. 2). Our phylogenetic analysis also showed that the species that feature in most evolutionary and ecological studies (*I. elegans*, *I. graellsii*, *I. ramburii* and *I. senegalensis*) are part of a well-supported lineage (*I. senegalensis* to *I. rufostigma;* Fig. 2), with a common ancestor less than 10 mya (Fig. 2). We found that all species from Australia and the South Pacific islands, except *I. heterosticta,* form a single clade *(I. taitensis* to *I. armstrongi*; Fig. 2). Similarly, we recovered a strongly supported Palearctic clade *(I. ezoin* to *I. intermedia;* Fig. 2) and strongly supported lineages distributed primarily in the Neotropical (*I. prognata* to *I. cruzi*) and Nearctic (*I. verticalis* to *I. kellicotti*) regions (Fig. 2).

**Figure 2.**
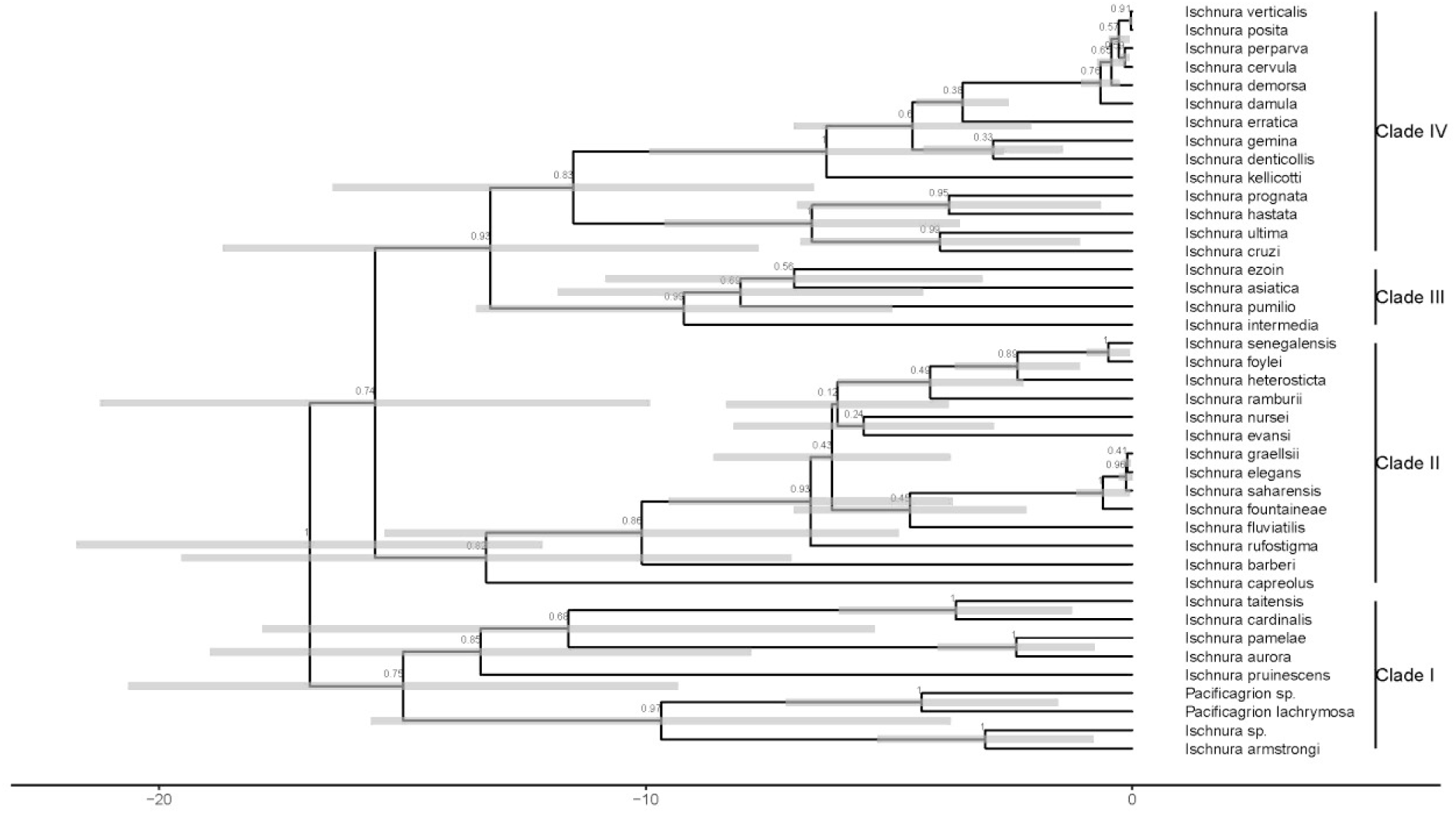
Time calibrated phylogeny of 41 species of damselflies from the genus *Ischnura* as estimated in StarBEAST2. Time axis represents millions of years from the present and grey bars at nodes indicate 95% HPD intervals for age estimates. Values at nodes indicate posterior probability of internal nodes.

**Figure 3.**
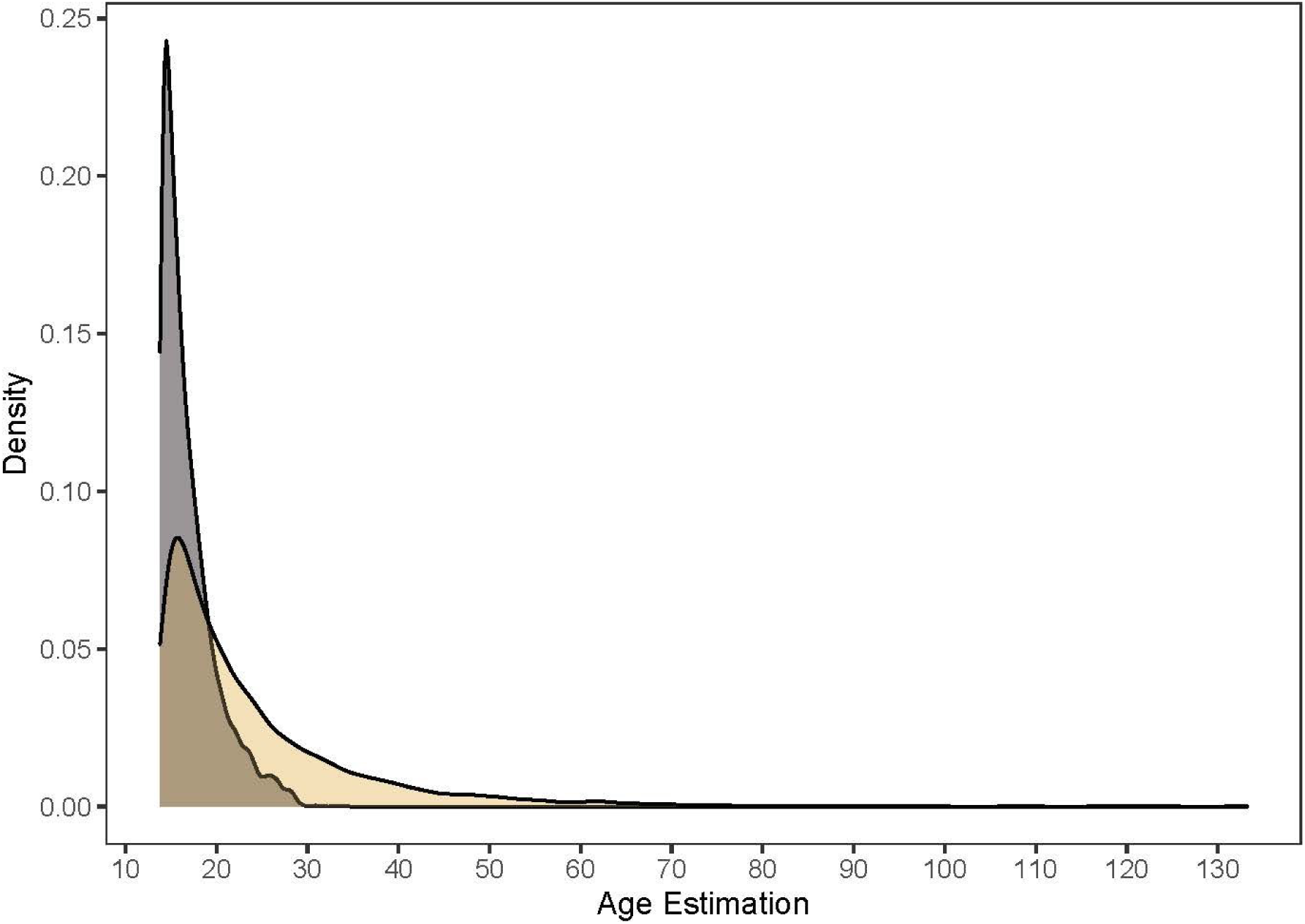
Posterior divergence time estimation at the root node sampling from the prior only (gold) and sampling with data (grey). Age estimation followed an exponentially distributed prior with an offset of 13.8 my to prevent improbable age estimation (see Methods).

### 2.2 Ancestral state reconstruction of female colour polymorphism

Out of the 41 species included in this study, three were FM(A), 13 were FM(H), 22 were FP(D), and three were FP(T) (cf. Fig 1B with Fig. 4). The MRCA of *Ischnura* was reconstructed as being FM(H), i.e. the ancestor was sexually dimorphic, with a mean posterior probability (PP) of 81% (Fig. 4). In contrast, the probability that the ancestor was sexually monomorphic (i.e. androchrome females only, being similar to males) was only 7 %, strongly suggesting that male mimicry and FP are derived traits and not ancestral. The clade containing most taxa from the Pacific islands (Fig. 2) was also inferred as being ancestrally FM(H) (PP = 81%; Fig. 4). All FP(T) species shared an FP(T) common ancestor (PP = 100%; Fig. 4), most likely derived from an FP(D) state (Fig. 4). The results presented here contrast strikingly with the conclusions of a recent study, in which the MRCA of *Ischnura* was reconstructed as female-polymorphic with 63% probability (Sánchez-Guillén et al., 2020). Repeating the analysis above but with the inclusion of nine additional species from Sánchez-Guillén et al. (2021) that were not included in our original taxa did not qualitatively alter our conclusion that the ancestor of the genus *Ischnura* was instead FM (H) (Fig. S2), although the support for a FM (H) ancestral state in *Ischnura* decreased to 66 % with the additional taxa (Supporting Results).

**Figure 4.**
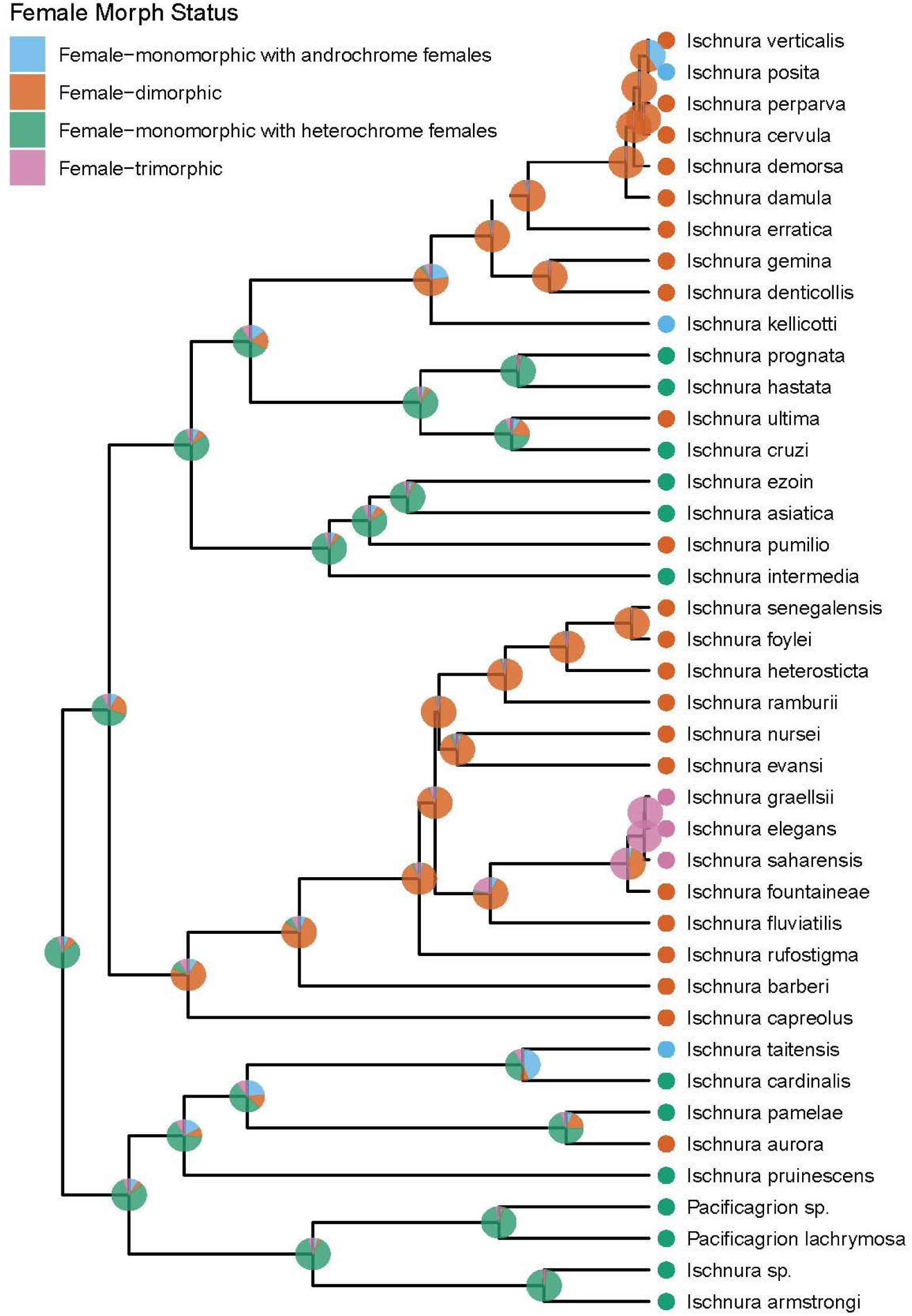
Ancestral state reconstruction across the *Ischnura* phylogeny. Extant morph states are represented by circles at tips and reconstructed ancestral states are represented by piecharts at nodes (female-heterochromatic only (FM(H)), green; female-andromorphic only (FM(A)), blue; female-dimorphic (FP(D)), red; female-trimorphic (FP(T)), pink). Piecharts show the mean posterior probability for each morph state and are plotted on the MCC tree.

Two evolutionary transitions occurred with high posterior probability during the diversification of *Ischnura*, according to the RJ MCMC analysis. The transition rate from FP(D) to FM(A) (i.e. the evolutionary loss of the heterochrome female morph) was above zero in 99.7% of the posterior (Fig. S1). Similarly, the transition from FM(H) to FP(D) (i.e. the evolutionary gain of an androchrome female morph) occurred with non-zero rate in 94.2% of the posterior (Fig. S1).

In contrast, the least probable transition, from FM(H) to FP(T) occurred in only 6.7% of the posterior (Fig. S1). The data on female colour states of extant *Ischnura* species and the posterior distribution of *Ischnura* time trees are therefore consistent with multiple evolutionary histories and different combinations of other character-state transitions being present or absent.

### 3.3 Geographic range size and female-colour polymorphism

Current geographic range sizes varied from 4.3 x 10^7^ km^2^ in the largely subtropical species *I. senegalensis*, which is distributed from sub-Saharan Africa to Japan, to 222 km^2^ to species endemic to American Samoa (Table S2). We found that all island-endemic species are FM, except for *I. foylei*, which is FP(D) and only known from the Jambi province in Sumatra, Indonesia (Table S2). In contrast, the three species distributed across the longitudinal range of the Old World, *I. senegalensis*, *I. pumilio* and *I. elegans,* are either FP(D) or FP(T). The probability of a species being FP increased significantly with its geographic range size (PMCMC = 0.003; Fig. 5). The probability of female colour polymorphism increased by ∼ 5% (posterior mean = 0.051, 95% HPD interval = 0.008 – 0.990) with an increase of 1 x10^6^ km^2^ in the geographic range size.

**4.** Discussion

**Figure 5.**
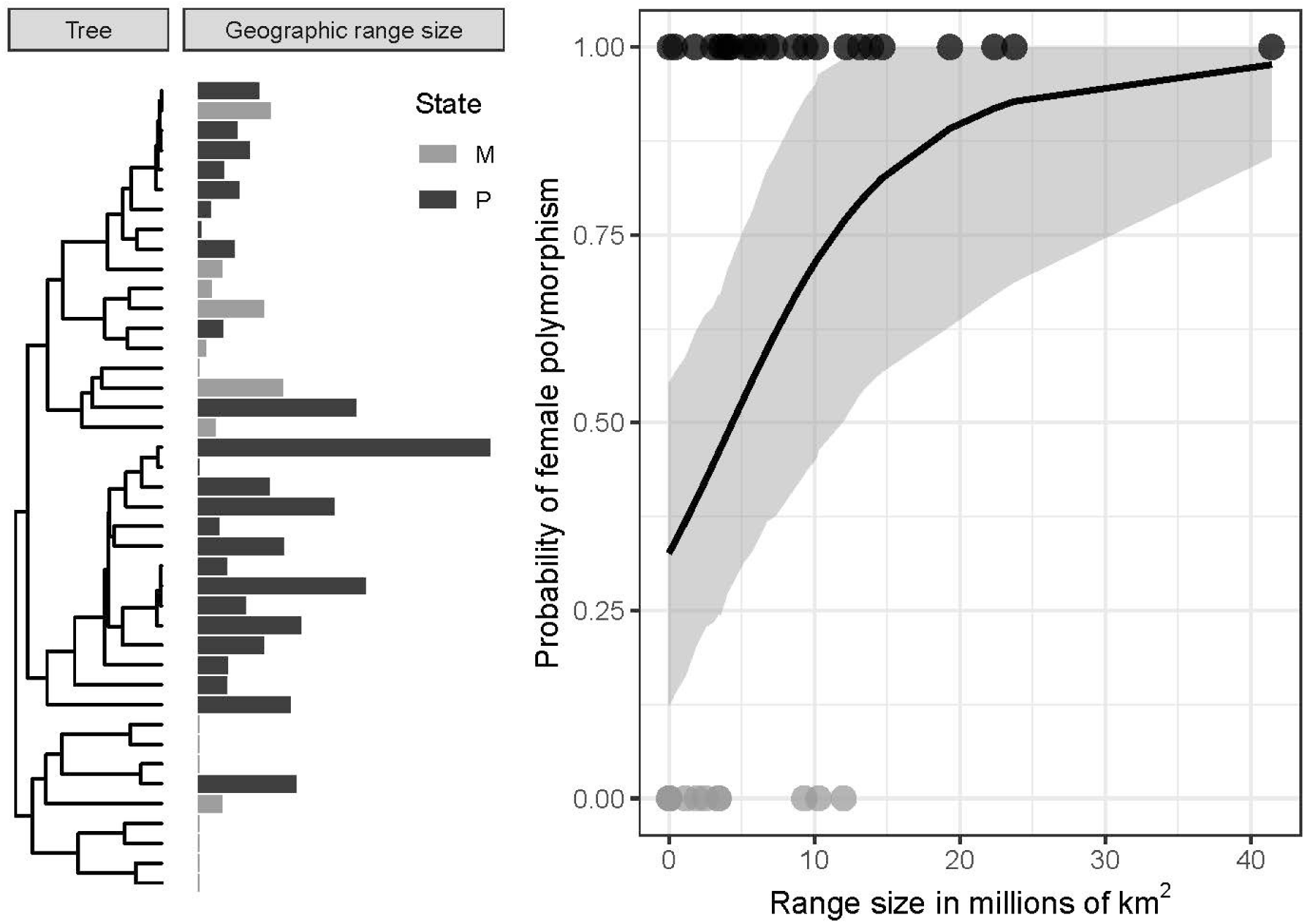
Probability of species having female polymorphism in relation to the species geographic range size, estimated in millions of squared kilometres. The left panel shows the MMC tree for the genus *Ischnura*. Bars on the centre panel are proportional to the geographic range of each species, except for species with very small ranges (<10^5^ Km^2^) which are all depicted by a similarly thin line. Light grey bars represent female-monomorphic (M) species and dark grey bars correspond to female-polymorphic (P) species. On the right panel, each circle represents an extant species. The black line shows the prediction from a Bayesian phylogenetic mixed-effect model and the grey area shows the 95% credible interval of the prediction. Species with larger geographic ranges are more likely to have polymorphic females (see Results).

Here, we have presented the first molecular time-calibrated phylogeny for the damselfly genus *Ischnura*. Although both dragonflies (suborder Anisoptera) and damselflies (suborder Zygoptera) have long been popular study organisms in ecology, evolution and conservation biology (Cordoba-Aguilar, 2008), time-calibrated molecular phylogenies that could facilitate phylogenetic comparative studies in these two insect groups are still relatively few (Callahan and McPeek, 2016; Damm et al., 2010; Letsch et al., 2016; Toussaint et al., 2019; Waller and Svensson, 2017). Several species in the genus *Ischnura* in particular have been subject to much research on sexual selection, sexual conflict and female colour polymorphism (see Introduction). The general branching patterns in this new phylogeny are largely and qualitatively in agreement with previous molecular phylogenies covering a subset of the taxa included here (Chippindale et al., 1999; Willink, 2018; Willink et al., 2019).

Using a UCLN molecular clock and fossil calibration, we inferred the MRCA of *Ischnura* as living between 13.8 and 23.4 mya, that is, the early Miocene. This is earlier than the MRCA of the closely related damselfly genus *Enallagma*, which has an estimated age of about 10 my (mid- to late Miocene), and where many speciation events seem to have been driven by glacial cycles during the Pleistocene (Callahan and McPeek, 2016). An early Miocene origin of *Ischnura* is also slightly earlier than the inferred origin of the closely related Palearctic damselfly genus *Coenagrion,* during the latter part of Miocene 15 mya (Swaegers et al., 2014), and the two genera *Melanosobasis* and *Nesobasis* in Fiji originating during late-middle Miocence (8-12 mya)(Beatty et al., 2017). We emphasize that our age estimate for the MRCA of *Ischnura* rests on the assumption that *I. velteni* is a crown fossil (Bechly, 2000). Future studies based on alternative dating strategies will be important to validate this inferred early Miocene origin of *Ischnura*.

According to our phylogeny and ancestral state reconstruction, the four most well-studied species in *Ischnura* (*I. senegalensis*, *I. graellsii, I. elegans* and *I. ramburii*)(Gering, 2017; Robertson, 1985; Sirot et al., 2003; Sirot and Brockmann, 2001) are part of a single clade, likely descended from an FP(D) ancestor (Fig. 4), that lived less than 10 mya (Figs. 2,4). While the genus *Ischnura* has increasingly attracted attention in ecological and microevolutionary studies, there is much untapped potential in understanding its macroevolutionary history and repeated transitions between female colour states. Our ancestral state reconstruction suggests that the ancestor of the genus *Ischnura* was FM(H), although we caution that the support for this inference was 81% (Fig. 4). Such a sexually dimorphic ancestor would imply multiple evolutionary transitions to obtain female-dimorphism, through the evolutionary emergence of a male-mimic. This evolutionary transition occurred with a non-zero rate in 94.2% of the posterior distribution of transition rate estimates in our RJ MCMC analysis (Fig. S1). This suggests that female colour polymorphism, with heterochrome and androchrome females coexisting within species, has evolved convergently multiple times in this genus, presumably as an evolutionary response to increased intensity of sexual conflict. Specifically, the emergence of a novel male-mimic might have been possible if it gained an initial fitness benefit due to advantages in intersexual mimicry through reduced male mating harassment (Robertson, 1985; Svensson et al., 2009; Willink et al., 2019). Such male-mimics might subsequently have increased to high frequencies in several species where they persisted alongside with the ancestral heteromorph as stable trans-species polymorphisms (Jamie and Meier, 2020). Currently, these female-limited polymorphisms are most likely maintained by frequency- and/or density dependent sexual selection caused by male mating harassment (Gosden and Svensson, 2009; Le Rouzic et al., 2015; Svensson et al., 2005; Takahashi et al., 2014, 2010).

We further note that the only known FP(T) clade, containing the well-studied European species *elegans* and *I. graellsii,* most likely evolved from an FP(D) ancestor (Fig. 4). The evolutionary emergence of a third female morph in an already FP system might indicate that sexual conflict through male mating harassment was particularly intense in this clade, which would fit with available empirical evidence from previous field and experimental studies (Cordero, 1992; Cordero et al., 1998; Gosden and Svensson, 2009; Takahashi et al., 2014; Willink et al., 2019). In support of this, *I. graellsii* and *I. elegans* have some of the longest copulation durations of all investigated species so far and mate for several hours (Robinson and Allgeyer, 1996). The long copulation duration in these two FP(T) species contrasts with the much shorter copulations and lower mating rates of some of the North American FM species, some of which have also been suggested to be monandrous (Robinson and Allgeyer, 1996). Thus, high mating rates, high degree of density-dependent male mating harassment (Gosden and Svensson, 2009) and long copulation duration (Robinson and Allgeyer, 1996) in the FP(T) clade containing *I. elegans* and *I. graellsii* might jointly have favoured the evolutionary establishment of a third morph in these species. Theory suggest that one evolutionary outcome of sexual conflict is the emergence of multiple genetic clusters in both males and females (Gavrilets and Waxman, 2002; Hayashi et al., 2007). There is small but growing body of empirical evidence that female phenotypic polymorphisms might have evolved as a response to sexual conflict in several different taxa (Iversen et al., 2019; Karlsson et al., 2013; Lee et al., 2019; Moon and Kamath, 2019; Reinhardt et al., 2007; Svensson et al., 2005).

The results in the present study run counter to to the conclusions of two previous phylogenetic comparative studies where the MRCA of *Ischnura* was reconstructed as FP (Mattern and Van Gossum, 2008; Sánchez-Guillén et al., 2020). Sánchez-Guillén et al. (2020) used a model of trait evolution, but assumed a single rate of trait evolution and only two possible character states (FM or FP), ignoring variation in sexual dimorphism among FM species and morph number (i. two or three morphs) in the FP taxa. In contrast, here we allowed for model uncertainty by using a reversible jump MCMC algorithm that samples alternative trait evolution models in proportion to their posterior probability (Pagel and Meade, 2006). Moreover, unlike Sánchez-Guillén et al. (2020), our ancestral state inference in the present study explicitly incorporated phylogenetic uncertainty, which through the MSC, captured uncertainty in how gene genealogical histories and species genealogical relate to one another.

To assess the possibility of disparate taxonomic sampling causing this discrepancy between studies, we included the unique species in Sánchez-Guillén et al. (2020) in a new posterior sample of species-trees, taking uncertainty in phylogenetic placement into consideration. We obtained qualitatively identical results as in our original analysis (Fig. S2). The newly added taxa from Sánchez-Guillén et al. (2020) were spread across one of the two lineages splitting from the MRCA of *Ischnura* in the present study (Fig. S2). The other early-splitting clade, however, included primarily South Pacific taxa, most of which are island-endemics and FM, and with one exception, are absent from Sánchez-Guillén et al. (2020). Our more extensive taxon sampling of this primarily FM clade likely contributed to inferring a FM ancestral state for *Ischnura* in the present study. It is noteworthy that of the 23 described and remaining species in this genus that were missing in both Sánchez-Guillén et al. (2020) and the present study, ten are endemic to small islands in the Pacific Ocean (Samoan islands, French Polynesian islands, Northern Mariana islands), one is endemic to Mauritius and at least six are highland endemics (Table S3). These relatively isolated taxa with restricted geographic ranges are more likely to be FM (Fig. 5; see below), and are likely to strengthen the hypothesis of ancestral female monomorphism in *Ischnura*, if they would become included in future phylogenetic studies of this genus.

Polymorphisms might promote speciation if different morphs diverge phenotypically in sympatry, followed by selective or random loss in allopatry, resulting in the fixation of alternative morphs in different populations (Jamie and Meier, 2020; West-Eberhard, 1986). This process – termed “morphic speciation” – predicts that monomorphic taxa are commonly found at the terminal tips on a phylogeny (Corl et al., 2010; West-Eberhard, 1986). There is some evidence for such a process among populations of colour polymorphic side-blotched lizards (*Uta stansburiana*)(Corl et al., 2010). Our results do not lend support for widespread morphic speciation, but are consistent with losses of heterochrome females in some FP(D) taxa, as this evolutionary transition occurred in 99.7% of the posterior in the RJ MCMC analysis. The most likely candidate for such potential morphic speciation is the common ancestor between *I. verticalis* and *I. posita* (Fig. 2). These two taxa are part of a North American clade that diversified rapidly during the Pleistocene (*I. damula* to *I. cervula;* Fig. 2). Such rapid speciation and diversification could have resulted from repeated glacial cycles, as in the closely related genus *Enallagma* that contains many sympatric young species in North America (Callahan and McPeek, 2016; McPeek and Gavrilets, 2006). Such glacial cycles and repeated Pleistocene bottlenecks could have led to morph fixation of a male-like female androchrome morph in the ancestor of *I. posita* during a period of allopatry (Fig. 4). *I. posita* is currently a widely distributed species, ranging from Nova Scotia in Canada to Central America (Paulson, 2012; Table S2). If morph loss in the ancestor of *I. posita* resulted from genetic drift during a period of allopatry and population contraction, it must then have been followed by rapid and recent range expansion. While this hypothesis remains to be investigated for *I. posita,* closely related and young sympatric taxa in *Enallagma* are thought to be the result of rapid speciation and range expansion after glacial retreat in the Quaternary (McPeek and Gavrilets, 2006; Turgeon et al., 2005).

Another potential example of morphic speciation leading to fixation of a single female morph is *I. kellicotti*, although it is less clear whether the lineage leading to *I. kellicotti* had an FP ancestor (Fig. 4). Nevertheless, the potential for morphic speciation in *I. kellicotti* is interesting because females display a striking pattern of ontogenetic colour changes. Sexually immature females start with a bright orange colouration that turns blue and male-like in mature females, and develops increasing pruinescence with age thereafter (Paulson, 2012, 2009). If used as a signal of reproductive availability (cf. Willink et al. 2019), the evolution of distinct developmental colour phases may have reduced male mating harassment and facilitated morphic speciation. We note that in several relatively distantly related species of *Ischnura*, immature females express orange or yellow colouration prior to reproductive maturity (e. g. in *ramburii, I. pumilio*, *I. senegalensis* and *I. capreolus*; Fig. 1E-G), and such striking immature colouration could signal reproductive unsuitability (cf. Willink et al. 2019). Alternatively, an evolutionary transition in habitat and mating system could have led to morph loss and morphic speciation in the ancestor of *I. kellicotti*. Individuals of *I. kelicotti* (“Lilypad forktail”) are typically found sitting on lilypad leaves in ponds (Paulson, 2012, 2009), consistent with a switch from non-territorial male-male scrambling competition over females with high mating harassment, to a new mating system of partial male territoriality with a lower degree of mating harassment. Such relaxed pressure from sexual conflict may have led to the loss of female polymorphism. A combination of “phylogenetic natural history” (Uyeda et al., 2018) and more detailed knowledge about the molecular and genetic basis of female colour development (Takahashi et al., 2018; Willink et al., 2020) could shed light on this issue and increase our general understanding of this and other macroevolutionary events in *Ischnura*.

The presence of female colour polymorphism was positively correlated with geographic range size (Fig. 5), which could come about in at least three different, but non-mutually exclusive, ways. First, species with large ranges are also likely to have large effective population sizes and this will decrease the effects of genetic drift and morph fixation. It is interesting to note here that several of the FM taxa are endemic to small islands in the Pacific Ocean (Table S2). Second, species with larger ranges and associated larger population sizes might experience stronger sexual conflict due to higher population densities, consistent with more intense male mating harassment under such conditions (Gosden and Svensson, 2009). Finally, FP species might also be able to exploit a broader range of niches and thereby also achieve higher population mean fitness compared to FM species, with positive effects on both population size and geographic range size (Forsman, 2016; Svensson, 2017; Takahashi et al., 2014; Takahashi et al., 2018; Takahashi and Noriyuki, 2019; West-Eberhard, 1986). Although the data in this study does not allow us to firmly distinguish between these non-mutually exclusive explanations for the positive relationship between the incidence of colour polymorphism and geographic range size (Fig. 5), these different scenarios deserve attention in future studies. A positive relationship between geographic range size and presence of colour polymorphism in *Ischnura* and other related damselflies was reported in a recent study, although without considering phylogenetic relatedness among species, as no molecular phylogeny was available at that time for *Ischnura* (Takahashi and Noriyuki, 2019).

Interestingly, the least supported nodes of our MCC tree are in the lineage from *I. senegalensis* to *I. rufostigma* (Fig. 2) and in the North American clade (*I. kellicotti* to *I. verticalis*; Fig. 2), which are both dominated by FP taxa (Fig. 4). These subsets of taxa were not particularly underrepresented in our dataset (Table S1). There are at least two potential biological explanations for this pattern. First, interspecific reproductive interference may be boosted by sexual conflict over mating rates (McPeek and Gavrilets, 2006; Svensson et al., 2009) which is likely to be more intense in FP species. This could result in more frequent hybridization and introgression in FP clades (Okude et al., 2020; Sanchez-Guillen et al., 2005; Sánchez-Guillén et al., 2011). Second, faster speciation when population sizes are large can result in deep coalescence among lineages and thus increase gene tree discordance (Maddison, 1997). If FP species have larger population sizes as our results suggest (Fig. 5), and colour polymorphisms accelerate speciation (Hugall and Stuart-Fox, 2012), then the speciation dynamics of FP clades may render them particularly challenging for phylogenetic inference. These alternative evolutionary scenarios are clearly worthy of further research, and require methods of phylogenetic inference such as the MSC, which accommodates incomplete lineage sorting and can be expanded to incorporate other biological processes such as introgression (Wen and Nakhleh, 2018; Zhang et al., 2018).

Finally, in one, and potentially two, of the FP clades, the female colour polymorphism seems to have arisen early and been subsequently maintained across multiple speciation events (Fig. 4). Therefore, the *Ischnura* female colour polymorphisms are examples of trans-species polymorphisms (Jamie and Meier, 2020), when multiple allelic variants become established in a population prior to speciation and are subsequently passed on to descendant taxa across multiple speciation events (Klein, 1987). Other examples of trans-species polymorphisms are rare, the most well-known being immune system polymorphisms and blood group variation in great apes (Azevedo et al., 2015) and self-incompatibility allelic systems in plants (Goldberg et al., 2010; Richman et al., 1996; Richman and Kohn, 2000). However, such trans-species polymorphisms may be more common than the limited current evidence suggests, particularly in colour polymorphic taxa (Corl et al., 2010; Jamie and Meier, 2020). In the damselfly genus *Ischnura*, some FP species have higher population-level fitness when female morph frequencies are even, compared to when they are biased towards any particular morph (Takahashi et al., 2014). In this study, we also showed that FP species have larger range sizes than FM species, indicating higher “ecological success” of the former (Fig. 5). Although these female colour polymorphisms are maintained by negative frequency-dependent selection at the individual level (Le Rouzic et al., 2015; Svensson et al., 2005; Takahashi et al., 2010) the higher-level population effects mentioned above might also indicate an additional role for species-level selection through biased extinction of FM taxa. Such higher level-effects would be similar to how self-incompatibility alleles are maintained in plant mating systems by species selection (Goldberg et al., 2010).

## Conclusions

The results in this study suggest that the damselfly genus *Ischnura* originated in the early Miocene, with ancestral females being monomorphic and markedly distinct from males in their colour pattern (heterochrome). The female colour polymorphisms in this genus arose on several occasions through repeated origins of a male-mimic in different species. Our ancestral state reconstruction further suggests that female colour polymorphisms have survived through multiple speciation events in at least two clades, contrary to the expectation of speciation by selective or random morph loss in allopatric populations (morphic speciation). However, we find at least one potential loss of the female colour polymorphism that could have resulted in morphic speciation. The multiple transitions in female-morph states across the phylogeny imply that the selective pressures from sexual conflict differ between taxa, and that the strength of frequency-dependent selection might be related to local population sizes and densities, with consequences for geographic range size and potentially also for extinction risk. This study provides a foundation for future micro- and macroevolutionary studies on how selective and neutral processes have shaped the evolution of discrete female colour morphs in *Ischnura* damselflies, by providing a phylogeny for future comparative studies in this genus.

## Acknowledgements

We are grateful to Debora Goedert and Masahito Tsuboi for constructive criticisms of an early draft of this manuscript. We wish to thank Tammy Ho, Chiara De Pasqual and Eva Friman for assistance in obtaining molecular sequence data. We also wish to thank all the following odonatologists who kindly provided samples for the phylogenetic analyses and shared their observations on colour variation in *Ischnura* species: Sylvain Charlat, Peter and Elaine Cowan, Nathalia von Ellenrieder, Sonia Ferreira, Kawsar Khan, Oleg Kosterin, Viktor Nilsson-Örtman, Ryo Futahashi, Roselyn and David Sparrow, Oliver Flint, curator of Odonata in the National Entomological Collection at the Smithsonian Institute, and Bill Mauffray, curator of Odonata at the Florida State Collection of Arthropods. Special thanks to D. J. Paulson, whose two volumes on North American odonates were extremely useful in our phenotypic data collection. Specimens were collected with permits from the Guyana Environmental Protection Agency (Reference No. 010715 BR001 and 012615 SP: 002), from Secretaría de Ambiente y Desarrollo Sustentable, Argentina (Ref No. 11084/16) and from the Cameroon Ministry of Forestry and Wildlife (Ref. No. 0000034). We acknowledge the logistic support from the Iwokrama International Centre for Rain Forest Conservation and Development, Guyana, the Karanambu Trust, Guyana, and the National Park Administration, Argentina. Funding for this study was provided by research grants from The Swedish Research Council (VR: grant no. 2016-03356), Carl Tryggers Foundation (CTS), Gyllenstiernska Krapperupstiftelsen (grant no. KR2018-0038), Lunds Djurskyddsfond, The Royal Physiographic Society in Lund, Stiftelsen Olle Enqvist Byggmästare and Stina Werners Foundation to E.I.S. and from The Royal Physiographic Society of Lund, the Jörgen Lindströms Fund and “Lunds Djurskyddsfond” to B.W.

## Data Availability

Novel sequence data has been made available on the National Centre for Biotechnology Information (https://www.ncbi.nlm.nih.gov/genbank/). All raw data, command files and scripts are available at GitHub (https://github.com/rachelblow/IschnuraPolymorphismEvolutionaryHistory) and Dryad Digital Repository (https://datadryad.org/stash) including convergence and stationarity checks of MCMC analyses.

