## Supporting Material for "A molecular phylogeny of forktail damselflies (genus *Ischnura*) reveals a dynamic macroevolutionary history of female colour polymorphisms"

**Table. S1.** NCBI accession numbers of sequences used in this study to construct a time-calibrated phylogeny of *Ischnura* damselflies. Two mitochondrial (16S, COI) and three nuclear (D7, PMRT, H3) loci were used (See Methods). Missing sequences from specimens for which not all markers were sequenced are denoted with '--'. Samples sequenced for this study are marked in **bold**. Samples marked with * indicate representative sequences used for extemded phylogenetic analyses with PASTIS (see Supplementary Methods). Sequence data downloaded from NCBI GenBank come from published studies: 1 = Willink et al. (2019); 2 = Karube et al. (2012); 3 = Dijkstra et al. (2014), 4 = Bybee et al. (2008); 5 = Ferreira et al. (2014); 6 = Kim et al. (2014).

| Taxon | Sample ID | 16S | COI | D7 | PMRT | H3 |
|---|---|---|---|---|---|---|
| *Ischnura armstrongi** | BEA546[1] | MK874546 | MK818631 | MK874530 | MK818716 | MK818667 |
| *Ischnura armstrongi* | **BEA575** | MT678014 | MT680671 | MT677959 | MT665861 | MT665812 |
| *Ischnura asiatica* | **BEA276** | MT678017 | MT680684 | MT677961 | MT665863 | MT665815 |
| *Ischnura asiatica** | BEA493[1] | MK874567 | MK818665 | MK874540 | MK818725 | MK818672 |
| *Ischnura asiatica* | **BEA526** | MT678018 | MT680685 | MT677962 | MT665864 | MW556203 |
| *Ischnura asiatica* | Kar1[2] | AB707553 | AB708497 | -- | -- | -- |
| *Ischnura aurora* | **BEA528** | MT678020 | MT680687 | MT677963 | -- | -- |
| *Ischnura aurora** | BEA548[1] | MK874550 | MK818649 | MK874533 | MK818720 | MK818673 |
| *Ischnura aurora* | **BEA549** | MT678021 | MT680690 | MT677964 | -- | MT665817 |
| *Ischnura aurora* | **BEA550** | MT678022 | MT680689 | MT677965 | -- | MT665818 |
| *Ischnura aurora* | **BEA685** | MT678023 | MT680688.2 | MT677966 | -- | MT665819 |
| *Ischnura aurora* | **BEA686** | MT678024 | MT680691 | MT677967 | MT665865 | MT665820 |
| *Ischnura aurora* | Dij1[3] | KF369749 | KF369414 | KF370148 | -- | -- |
| *Ischnura aurora* | **BEA123** | MT678019 | MT680686 | -- | -- | MT665816 |
| *Ischnura aurora* | Kar1[2] | AB707554 | AB708498 | -- | -- | -- |
| *Ischnura aurora* | Kar2[2] | AB707555 | AB708499 | -- | -- | -- |
| *Ischnura aurora* | Kar3[2] | AB707556 | AB708500 | -- | -- | -- |
| *Ischnura aurora* | Kar4[2] | AB707557 | AB708501 | -- | -- | -- |
| *Ischnura aurora* | Kar5[2] | AB707558 | AB708502 | -- | -- | -- |
| *Ischnura aurora* | Kar6[2] | AB707559 | AB708503 | -- | -- | -- |
| *Ischnura barberi** | BEA104[1] | MK874570 | MK818637 | MK874521 | MK818714 | MK818674 |
| *Ischnura barberi* | Byb1[4] | -- | -- | EU055231 | -- | EU055427 |
| *Ischnura capreolus* | **BEA010** | MT678025 | MT680676 | MT677968 | -- | -- |
| *Ischnura capreolus** | BEA411[1] | MK874555 | MK818636 | MK874536 | MK818724 | MK818675 |

| Taxon | Sample ID | 16S | COI | D7 | PMRT | H3 |
|---|---|---|---|---|---|---|
| *Ischnura capreolus* | **BEA641** | MT678026 | MT680678 | MT677969 | MT665866 | MT665821 |
| *Ischnura capreolus* | **BEA727** | MT678027 | MT680677 | MT677970 | -- | MT665822 |
| *Ischnura capreolus* | **BEA742** | MT678028 | MT680679 | MT677971 | MT665867 | MT665823 |
| *Ischnura cardinalis** | BEA675[1] | MK874552 | MK818633 | MK874534 | MK818722 | MK818676 |
| *Ischnura cardinalis* | **BEA688** | MT678029 | MT680674 | MT677972 | -- | MT665824 |
| *Ischnura cervula** | BEA449[1] | MK874556 | MK818653 | MK874512 | MK818728 | MK818677 |
| *Ischnura cervula* | **BEA450** | MT678030 | MT680699 | MT677973 | MT665868 | MT665825 |
| *Ischnura cervula* | Fer1[5] | -- | -- | -- | KM276622 | -- |
| *Ischnura cruzi** | BEA078[1] | MK874582 | MK818661 | MK874523 | MK818734 | MK818669 |
| *Ischnura damula** | BEA087[1] | MK874560 | MK818655 | MK874509 | MK818732 | MK818678 |
| *Ischnura demorsa** | BEA075[1] | MK874561 | MK818656 | MK874541 | MK818729 | MK818679 |
| *Ischnura denticollis** | BEA217[1] | -- | -- | -- | MK818738 | MK818680 |
| *Ischnura denticollis* | Fer1[5] | -- | -- | -- | KM276603 | -- |
| *Ischnura elegans** | BEA494[1] | MK874571 | MK818645 | MK874517 | MK818711 | MK818681 |
| *Ischnura elegans* | **BEA682** | MT678031 | MT680720 | MT677974 | MT665869 | MT665826 |
| *Ischnura elegans* | **BEA689** | MT678032 | MT680721 | MT677975 | MT665870 | MT665827 |
| *Ischnura elegans* | **BEA690** | MT678033 | MT680722 | MT677976 | -- | MT665828 |
| *Ischnura elegans* | Dij1[3] | -- | -- | KF370149 | -- | -- |
| *Ischnura elegans* | Kar1[2] | AB707560 | AB708504 | -- | -- | -- |
| *Ischnura elegans* | Kar2[2] | AB707561 | AB708505 | -- | -- | -- |
| *Ischnura elegans* | Kar3[2] | AB707562 | AB708506 | -- | -- | -- |
| *Ischnura elegans* | Kim1[6] | KF256901 | KF257118 | -- | -- | -- |
| *Ischnura erratica** | BEA326[1] | -- | -- | MK874545 | -- | MK818682 |
| *Ischnura evansi** | BEA267[1] | MK874578 | MK818644 | MK874528 | -- | MK818683 |
| *Ischnura ezoin** | BEA527[1] | MK874568 | MK818666 | MK874537 | MW556205 | MW556204 |
| *Ischnura ezoin* | Kar1[2] | AB707523 | AB708467 | -- | -- | -- |
| *Ischnura ezoin* | Kar2[2] | AB707524 | AB708468 | -- | -- | -- |
| *Ischnura ezoin* | Kar3[2] | AB707525 | AB708469 | -- | -- | -- |
| *Ischnura ezoin* | Kar4[2] | AB707526 | AB708470 | -- | -- | -- |

| Taxon | Sample ID | 16S | COI | D7 | PMRT | H3 |
|---|---|---|---|---|---|---|
| *Ischnura ezoin* | Kar5[2] | AB707527 | AB708471 | -- | -- | -- |
| *Ischnura fluviatilis** | BEA043[1] | MK874579 | MK818640 | MK874519 | MK818709 | MK818684 |
| *Ischnura fluviatilis* | **BEA056** | MT678034 | MT680707 | MT677977 | MT665871 | MT665829 |
| *Ischnura fluviatilis* | **BEA643** | MT678036 | MT680709 | MT677978 | MT665872 | MT665831 |
| *Ischnura fluviatilis* | **BEA738** | MT678037 | MT680710 | MT677979 | MT665873 | MT665832 |
| *Ischnura fluviatilis* | **BEA642** | MT678035 | MT680708 | -- | -- | MT665830 |
| *Ischnura fountaineae** | BEA479[1] | MK874572 | MK818646 | MK874516 | MK818715 | MK818685 |
| *Ischnura foylei** | BEA673[1] | MK874575 | MK818642 | MK874532 | MK818705 | MK818686 |
| *Ischnura gemina** | BEA344[1] | MK874559 | MK818652 | MK874542 | MK818739 | MK818687 |
| *Ischnura graellsii* | **BEA425** | MT678038 | MT680723 | MT677980 | MT665874 | MT665833 |
| *Ischnura graellsii** | BEA694[1] | MK874573 | MK818647 | MK874518 | MK818713 | MK818688 |
| *Ischnura graellsii* | **BEA695** | MT678039 | MT680724 | MT677981 | -- | MT665834 |
| *Ischnura graellsii* | **BEA696** | MT678040 | MT680725 | MT677982 | -- | MT665835 |
| *Ischnura hastata* | **BEA051** | MT678041 | MT680702 | MT677983 | MT665875 | MT665836 |
| *Ischnura hastata** | BEA705[1] | MK874584 | MK818660 | MK874526 | MK818736 | MK818689 |
| *Ischnura hastata* | **BEA706** | MT678042 | MT680703 | MT677984 | MT665876 | MT665837 |
| *Ischnura hastata* | Fer1[5] | -- | -- | -- | KM276605 | -- |
| *Ischnura hastata* | Fer1[5] | -- | -- | -- | KM276606 | -- |
| *Ischnura hastata* | Fer1[5] | -- | -- | -- | KM276607 | -- |
| *Ischnura hastata* | Fer1[5] | -- | -- | -- | KM276608 | -- |
| *Ischnura hastata* | Fer1[5] | -- | -- | -- | KM276609 | -- |
| *Ischnura hastata* | Fer1[5] | -- | -- | -- | KM276610 | -- |
| *Ischnura hastata* | Fer1[5] | -- | -- | -- | KM276611 | -- |
| *Ischnura hastata* | Fer1[5] | -- | -- | -- | KM276612 | -- |
| *Ischnura hastata* | Fer1[5] | -- | -- | -- | KM276613 | -- |
| *Ischnura hastata* | Fer1[5] | -- | -- | -- | KM276614 | -- |
| *Ischnura hastata* | Fer1[5] | -- | -- | -- | KM276615 | -- |
| *Ischnura hastata* | Fer1[5] | -- | -- | -- | KM276616 | -- |
| *Ischnura hastata* | Fer1[5] | -- | -- | -- | KM276617 | -- |

| Taxon | Sample ID | 16S | COI | D7 | PMRT | H3 |
|---|---|---|---|---|---|---|
| *Ischnura hastata* | Fer1[5] | -- | -- | -- | KM276618 | -- |
| *Ischnura hastata* | Fer1[5] | -- | -- | -- | KM276619 | -- |
| *Ischnura hastata* | Fer1[5] | -- | -- | -- | KM276620 | -- |
| *Ischnura heterosticta** | BEA551[1] | MK874577 | MK818641 | MK874513 | MK818707 | MK818690 |
| *Ischnura heterosticta* | **BEA552** | MT678044 | MT680712 | MT677985 | MT665877 | MT665838 |
| *Ischnura heterosticta* | **BEA122** | MT678043 | MT680711 | -- | -- | -- |
| *Ischnura heterosticta* | Kar1[2] | AB707563 | AB708507 | -- | -- | |
| *Ischnura intermedia** | BEA676[1] | MK874566 | MK818663 | MK874539 | MK818727 | MK818691 |
| *Ischnura intermedia* | **BEA691** | MT678045 | MT680680 | MT677986 | -- | MT665839 |
| *Ischnura kellicotti** | BEA105[1] | MK874565 | MK818651 | MK874515 | MK818740 | MK818692 |
| *Ischnura kellicotti* | Fer1[5] | -- | -- | -- | KM276621 | |
| *Ischnura nursei** | Dij1[3] | KF369893 | KF369538 | KF370292 | -- | -- |
| *Ischnura pamelae** | BEA451[1] | MK874551 | MK818650 | MK874535 | MK818721 | MK818693 |
| *Ischnura perparva** | BEA084[1] | MK874557 | MK818654 | MK874508 | MK818730 | MK818671 |
| *Ischnura posita** | **BEA083[1]** | MK874564 | MT680692 | MK874510 | MK818731 | MK818670 |
| *Ischnura posita* | **BEA708** | MT678046 | MT680697 | MT677987 | MT665878 | MT665840 |
| *Ischnura posita* | **BEA709** | MT678047 | MT680701 | MT677988 | -- | MT665841 |
| *Ischnura posita* | **BEA710** | MT678048 | MT680700 | MT677989 | -- | MT665842 |
| *Ischnura posita* | **BEA711[1]** | MT678049 | MK818657 | MT677990 | MT665879 | MT665843 |
| *Ischnura posita* | **BEA712** | -- | -- | MT677991 | MT665880 | MT665844 |
| *Ischnura posita* | Fer1[5] | -- | -- | -- | KM276604 | -- |
| *Ischnura prognata** | BEA239[1] | MK874583 | MK818659 | MK874511 | MK818737 | -- |
| *Ischnura pruinescens** | BEA353[1] | MK874554 | MK818635 | -- | -- | MK818694 |
| *Ischnura pumilio* | **BEA375** | MT678050 | MT680683 | MT677992 | MT665881 | MT665845 |
| *Ischnura pumilio* | **BEA525** | MT678051 | MT680681 | MT677993 | MT665882 | MT665846 |
| *Ischnura pumilio** | **BEA718[1]** | MK874569 | MK818664 | MK874527 | MT665883 | MK818695 |
| *Ischnura pumilio* | **BEA719[1]** | MT678052 | MT680682 | MT677994 | MK818726 | MT665847 |
| *Ischnura ramburii* | **BEA155** | -- | -- | MT677995 | MT665884 | MT665848 |
| *Ischnura ramburii** | BEA412[1] | MK874581 | MK818638 | MK874529 | MK818708 | MK818696 |

| Taxon | Sample ID | 16S | COI | D7 | PMRT | H3 |
|---|---|---|---|---|---|---|
| *Ischnura ramburii* | **BEA426** | MT678053 | MT680705 | MT677996 | MT665885 | MT665849 |
| *Ischnura rufostigma* | **BEA132** | MT678054 | MT680706 | MT677997 | -- | -- |
| *Ischnura rufostigma** | BEA465[1] | MK874580 | MK818639 | MK874538 | MK818710 | MK818697 |
| *Ischnura rufostigma* | Kar1[2] | AB707564 | AB708508 | -- | -- | -- |
| *Ischnura saharensis** | BEA481[1] | MK874574 | MK818648 | MK874525 | MK818712 | MK818698 |
| *Ischnura saharensis* | **BEA430** | MT678055 | MT680726 | -- | MT665886 | MT665850 |
| *Ischnura senegalensis* | **BEA474** | MT678056 | MT680714 | MT677998 | MT665887 | -- |
| *Ischnura senegalensis* | **BEA477** | MT678057 | MT680715 | MT677999 | MT665888 | -- |
| *Ischnura senegalensis* | **BEA478** | MT678058 | MT680716 | MT678000 | MT665889 | -- |
| *Ischnura senegalensis* | **BEA488** | MT678059 | MT680717 | MT678001 | MT665890 | -- |
| *Ischnura senegalensis* | **BEA490** | MT678060 | MT680718 | MT678002 | MT665891 | MT665851 |
| *Ischnura senegalensis* | **BEA491** | MT678061 | MT680719 | MT678003 | MT665892 | -- |
| *Ischnura senegalensis* | **BEA495** | MT678062 | MT680713 | MT678004 | MT665893 | MT665852 |
| *Ischnura senegalensis** | BEA511[1] | MK874576 | MK818644 | MK874524 | MK818706 | MK818699 |
| *Ischnura senegalensis* | Dij1[3] | -- | -- | KF370150 | -- | -- |
| *Ischnura senegalensis* | Kar1[2] | AB707565 | AB708509 | -- | -- | -- |
| *Ischnura senegalensis* | Kar2[2] | AB707566 | AB708510 | -- | -- | -- |
| *Ischnura senegalensis* | Kar3[2] | AB707567 | AB708511 | -- | -- | -- |
| *Ischnura senegalensis* | Kim1[6] | KF256888 | KF257106 | -- | -- | -- |
| *Ischnura* sp. | **BEA545[1]** | MT678015 | MK818632 | MT677960 | -- | MT665813 |
| *Ischnura* sp.* | **BEA670[1]** | MK874547 | MT680672 | MK874520 | MK818717 | MK818668 |
| *Ischnura* sp. | **BEA671** | MT678016 | MT680673 | -- | MT665862 | MT665814 |
| *Ischnura taitensis** | BEA674[1] | MK874553 | MK818634 | MK874531 | MK818723 | MK818700 |
| *Ischnura taitensis* | **BEA687** | MT678063 | MT680675 | MT678005 | MT665894 | MT665853 |
| *Ischnura ultima** | BEA554[1] | MK874585 | MK818662 | MK874522 | MK818735 | MK818701 |
| *Ischnura ultima* | **BEA553** | MT678064 | MT680704 | MT678006 | -- | MT665854 |
| *Ischnura verticalis** | **BEA111[1]** | MK874563 | MK818658 | MT678007 | MK818733 | MK818702 |
| *Ischnura verticalis* | **BEA713** | MT678065 | MT680698 | MT678008 | MT665895 | MT665855 |
| *Ischnura verticalis* | **BEA714** | MT678066 | MT680695 | MT678009 | MT665896 | MT665856 |

| Taxon | Sample ID | 16S | COI | D7 | PMRT | H3 |
|---|---|---|---|---|---|---|
| *Ischnura verticalis* | **BEA715** | MT678067 | MT680696 | MT678010 | MT665897 | MT665857 |
| *Ischnura verticalis* | **BEA716** | MT678068 | MT680693 | MT678011 | MT665898 | MT665858 |
| *Ischnura verticalis* | **BEA717**[1] | MT678069 | MT680694 | MK874514 | MT665899 | -- |
| *Pacificagrion lachrymosa* | **BEA560** | MT678070 | MT680670 | MT678012 | -- | MT665859 |
| *Pacificagrion lachrymosa** | BEA574[1] | MK874548 | MK818629 | MK874543 | MK818718 | MK818703 |
| *Pacificagrion* sp. | **BEA547** | MT678071 | MT680669 | MT678013 | -- | MT665860 |
| *Pacificagrion* sp.* | BEA576[1] | MK874549 | MK818630 | MK874544 | MK818719 | MK818704 |

**Table S2.** Character states and geographic range areas for 41 species of *Ischnura* damselflies included in this study. The variable 'State' indicates whether female-limited polymorphism (FP) occurs within a species, and if so, whether two (FP(D)) or three (FP(T)) female morphs are present, and in the case that a single morph exists, whether females are visually distinct from males in their colour pattern (heterochrome; FM(H)) or largely similar to males (androchrome; FM(A)). The variable 'StateBin' indicates if females are monomorphic (M), regardless of their colour pattern (heterochrome or androchrome), or if females are polymorphic (P), regardless of whether two or three female morphs co-occur. This was the response variable used in the geographic range size analysis (see Methods and Fig. 5). For each species, we report all countries of known occurrence, with names abbreviated according to the International Organization for Standardization (ISO). Whenever available we include lower-level administrative divisions, such as provinces, states and territories. For each species we calculate the total area of the geographic range by adding the areas of all administrative divisions where the species has been recorded, according to literature, online resources and museum collections listed under 'Sources'.

| Species | State | StateBin | Countries | Other divisions | Range area ($Km^2$) | Sources |
|---|---|---|---|---|---|---|
| *Ischnura armstrongi* | FM(H) | M | ASM, WSM | | 3074 | Marinov et al. (2015), Marinov (2020)* |
| *Ischnura asiatica* | FM(H) | M | CHN, HKG, JPN, KOR, PRK, RUS, TWN | Amur, Anhui, Beijing, Buryat, Chukot, Guangdong, Guangxi, Guizhou, Hebei, Heilongjiang, Henan, Hubei, Jiangsu, Jiangxi, Jilin, Kamchatka, Khabarovsk,Liaoning, Maga Buryatdan, Primor'ye, Sakha, Sakhalin, Shaanxi, Shandong, Shanghai, Sichuan,Tianjin, Xizang, Yevrey, Zabaykal'ye, Zhejiang | 12002217 | Haritonov and Malikova (1998), Ozono et al. (2012), Yu and Chen (2015), Seehausen and Fiebig (2016), Zhang et al. (2018), GBIF.org (2020), |
| *Ischnura aurora* | FP(D) | P | AUS, BGD, BTN, CHN, FJI, FSM, GUM, IDN, IND, JPN, LAO, LKA, MMR, MNP, NCL, NPL, PAK, PHL, PYF, SLB, TON, TWN, VUT, WSM | Andhra Pradesh, Arunachal Pradesh, Assam, Bihar, Fujian, Guangdong, Hainan, Himachal Pradesh, Kerala, Madhya Pradesh, Maharashtra, Manipur, Meghalaya, Nagaland, New South Wales, Northern Territory, Odisha, Ogasawara islands, Punjab, Queensland, Rajasthan, Ryukyu Islands, Sikkim, Tamil Nadu, Tasmania, Uttar Pradesh, Victoria, West Bengal, Western Australia | 13907704 | Rowe (1987), Ozono et al. (2012), Dow et al. (2020) |
| *Ischnura barberi* | FP(D) | P | MEX, USA | Arizona, California, New Mexico, Oklahoma, Texas, Utah | 4058740 | Paulson (2009) |
| *Ischnura capreolus* | FP(D) | P | ARG, BLZ, BRA, COL, CRI, ECU, SLV, GTM, GUF, GUY, HND, MEX, NIC, PAN, PER, PRY, SUR, TTO, VEN | Bahia, Espírito Santo, Mato Grosso, Pará, Pernambuco, Rio de Janeiro, Rio Grande do Sul, São Paulo | 13113468 | Heckman (2008), Kompier (2015), Vilela et al. (2017) |

| Species | State | StateBin | Countries | Other divisions | Range area (Km$^2$) | Sources |
|---|---|---|---|---|---|---|
| *Ischnura cardinalis* | FM(H) | M | PYF | | 3680 | Marinov et al. (2019) |
| *Ischnura cervula* | FP(D) | P | CAN, MEX, USA | Alberta, Arizona, British Columbia, California, Colorado, Idaho, Montana, New Mexico, Nevada, Oregon, Saskatchewan, Utah, Washington, Wyoming | 7288629 | Paulson (2009) |
| *Ischnura cruzi* | FM(H) | M | COL | | 1136512 | Heckman (2008), Realpe (2010) |
| *Ischnura damula* | FP(D) | P | CAN, USA | Alberta, British Columbia, Manitoba, Saskatchewan, Yukon, Arizona, Colorado, Nebraska, New Mexico, North Dakota, Texas, Utah, Wyoming | 5820275 | Paulson (2009) |
| *Ischnura demorsa* | FP(D) | P | MEX, USA | Arizona, Kansas, New Mexico, Oklahoma, Texas | 3642413 | Paulson (2009) |
| *Ischnura denticollis* | FP(D) | P | GTM, MEX, USA | Arizona, California, Idaho, Kansas, Nevada, New Mexico, Oklahoma, Oregon, Texas, Utah | 5135142 | Paulson (2009) |
| *Ischnura elegans* | FP(T) | P | ALB, AUT, BEL, BIH, BLR, BGR, CHE, CHN, CYP, CZE, DEU, DNK, ESP, EST, FIN, FRA, GBR, GRC, HRV, HUN, IDN, IND, IRL, IRN, ISR, ITA, JOR, JPN, KOR, LBN, LIE, LKA, LTU, LUX, LVA, MDA, MKD, MNE, MNG, MYS, NLD, NOR, NPL, PAK, PRK, POL, PSE, ROU, SRB, SVK, SVN, SWE, SYR, TUR, UKR | Himachal Pradesh, Jammu and Kashmir, Uttar Pradesh, West Bengal | 23744568 | Dijkstra and Lewington (2006), Dow (2010) |
| *Ischnura erratica* | FP(D) | P | CAN, USA | British Columbia, California, Oregon, Washington | 1784487 | Paulson (2009) |
| *Ischnura evansi* | FP(D) | P | AFG, ARE, BHR, EGY, IRN, IRQ, ISR, JOR, KAZ, KGZ, KWT, OMN, PSE, QAT, SAU, SDN, SYR, TJK, TKM, YEM | | 12210380 | Marzoq (2005), Clausnitzer et al. (2013)* |
| *Ischnura ezoin* | FM(H) | M | JPN | Ogasawara islands | 258 | Ozono et al. (2012), Karube et al. (2020) |
| *Ischnura fluviatilis* | FP(D) | P | ARG, BOL, BRA, CHL, ECU, GUF, GUY, PRY, URY, VEN | Mato Grosso, Pará, Rio de Janeiro, Rio Grande do Sul, São Paulo | 9375458 | Heckman (2008), von Ellenrieder |

| Species | State | StateBin | Countries | Other divisions | Range area ($Km^2$) | Sources |
|---|---|---|---|---|---|---|
| | | | | | | (2009), Kompier (2015) |
| *Ischnura fountaineae* | FP(D) | P | ARE, AZE, DZA, EGY, GEO, IRN, IRQ, ISR, JOR, KAZ, KGZ, LBR, LBY, MAR, OMN, PSE, QAT, RUS, SAU, SYR, TKM, TUN, TUR | Dagestan | 14656465 | Boudot et al. (2013), Schröter (2010), Seehausen et al. (2016) |
| *Ischnura foylei* | FP(D) | P | IDN | Jambi | 49095 | Kosterin (2015) |
| *Ischnura gemina* | FP(D) | P | USA | California | 409615 | Paulson (2009) |
| *Ischnura graellsii* | FP(T) | P | DZA, ESP, FRA, MAR, PRT, TUN | | 4023699 | Dijkstra and Lewington (2006), Clausnitzer (2009) |
| *Ischnura hastata* | FM(H) | M | BLZ, COL, CRI, CUB, DOM, SLV, GTM, GUY, HND, MEX, NIC, PAN, TTO, USA, VEN | Alabama, Arizona, Arkansas, Connecticut, Florida, Georgia, Illinois, Indiana, Kansas, Kentucky, Louisiana, Massachusetts, Michigan, Mississippi, Missouri, Nebraska, New Jersey, New Mexico, New York, North Carolina, Ohio, Oklahoma, Pennsylvania, Rhode Island, South Carolina, Tennessee, Texas, Virginia, West Virginia | 9301499 | Esquivel (2006), Heckman (2008), Paulson (2012) |
| *Ischnura heterosticta* | FP(D) | P | AUS, FJI, IDN, NCL, PLW, PNG, SLB, TON, VUT | New South Wales, Northern Territory, Queensland, South Australia, Tasmania, Victoria, Western Australia | 10113924 | Huang and Reinhard (2012), Dow (2017a) |
| *Ischnura intermedia* | FM(H) | M | CYP, IRN, SYR, TKM, TUR | Adana, Adiyaman, Antalya, Batman, Burdur, Diyarbakir, Gaziantep, Hatay, Isparta, K. Maras, Karaman, Kilis, Mardin, Mersin, Mugla, Osmaniye, Sanliurfa, Siirt, Sirnak | 2495078 | Knifj et al. (2016)* |
| *Ischnura kellicotti* | FM(A) | M | USA | Alabama, Arkansas, Connecticut, Delaware, District of Columbia, Florida, Georgia, Illinois, Indiana, Kentucky, Louisiana, Maine, Massachusetts, Michigan, Mississippi, Missouri, New Hampshire, New Jersey, New York, North Carolina, Oklahoma, Pennsylvania, Rhode Island, South Carolina, Tennessee, Texas, Vermont, Virginia, West Virginia | 3404330 | Paulson (2012) |

| Species | State | StateBin | Countries | Other divisions | Range area (Km$^2$) | Sources |
|---|---|---|---|---|---|---|
| *Ischnura nursei* | FP(D) | P | BGD, IND, PAK | Chandigarh, Chhattisgarh, Madhya Pradesh, Maharashtra, Odisha, Punjab, Rajasthan, Tamil Nadu, Uttar Pradesh, Uttarakhand, West Bengal | 3028389 | Subramanian (2010)[*] |
| *Ischnura pamelae* | FM(H) | M | NCL | | 18775 | Vick and Davies (1988), Kalkman (2020) |
| *Ischnura perparva* | FP(D) | P | | Alberta, British Columbia, California, Colorado, Idaho, Montana, Nebraska, Nevada, New Mexico, North Dakota, Oregon, Saskatchewan, South Dakota, Utah, Washington, Wyoming | 5624521 | Paulson (2009) |
| *Ischnura posita* | FM(A) | M | BLZ, CAN, GTM, MEX, USA | Alabama, Arkansas, Connecticut, Delaware, District of Columbia, Florida, Georgia, Illinois, Indiana, Iowa, Kansas, Kentucky, Louisiana, Maine, Manitoba, Massachusetts, Michigan, Minnesota, Mississippi, Missouri, Nebraska, New Hampshire, New Jersey, New York, Newfoundland and Labrador, North Carolina, Nova Scotia, Ohio, Oklahoma, Ontario, Québec, Pennsylvania, Rhode Island, South Carolina, Tennessee, Texas, Vermont, Virginia, Wisconsin, West Virginia | 10273446 | Paulson (2012) |
| *Ischnura prognata* | FM(H) | M | USA | Alabama, Florida, Georgia, Louisiana, Missouri, North Carolina, South Carolina, Tennessee, Texas, Virginia, West Virginia | 1915019 | Paulson (2012) |
| *Ischnura pruinescens* | FM(H) | M | AUS, IDN | Northern Territory, Papua, Queensland | 3392025 | Theischinger and Hawking (2006), Dow (2017b) |
| *Ischnura pumilio* | FP(D) | P | AFG, ALB, AND, ARM, AUT, AZE, BEL, BIH, BLR, BGR, CHE, CHN, CYP, CZE, DEU, DNK, ESP, EST, FIN, FRA, GBR, GRC, HRV, HUN, IRL, IRN, ISR, ITA, JOR, KAZ, KGZ, LBN, LIE, LTU, LUX, LVA, MAR, MDA, MKD, MNE, MNG, NLD, NOR, | Adygey, Arkhangel'sk, Astrakhan', Bashkortostan, Belgorod, Bryansk, Chechnya, Chelyabinsk, Chuvash, City of St. Petersburg, Dagestan, Ingush, Ivanovo, Kabardin-Balkar, Kaliningrad, Kalmyk, Kaluga, Karachay-Cherkess, Karelia, Khanty-Mansiy, Kirov, Komi, Kostroma, Krasnodar, Kurgan, Kursk, Leningrad, Lipetsk, Mariy-El, Mordovia, Moscow City, Moskva, Murmansk, Nei Mongol, Nenets, Nizhegorod, | 22384600 | Dijkstra and Lewington (2006), Boudot (2014) |

| Species | State | StateBin | Countries | Other divisions | Range area (Km$^2$) | Sources |
|---|---|---|---|---|---|---|
| | | | POL, PRT, ROU, RUS, SRB, SVK, SVN, SWE, SYR, TJK, TKM, TUN, TUR, UKR, UZB | North Ossetia, Novgorod, Orel, Orenburg, Penza, Perm', Pskov, Rostov, Ryazan', Samara, Saratov, Shanxi, Smolensk, Stavropol', Sverdlovsk, Tambov, Tatarstan, Tula, Tver', Tyumen', Udmurt, Ul'yanovsk, Vladimir, Volgograd, Vologda, Voronezh, Yamal-Nenets, Yaroslavl' | | |
| *Ischnura ramburii* | FP(D) | P | BHS, BLZ, BRA, CHL, COL, CRI, CUB, CYM, DOM, ECU, SLV, GLP, GTM, GUF, HND, HTI, JAM, MEX, MTQ, NIC, PAN, PER, PRI, PRY, SUR, TTO, VEN | Alabama, Arizona, Arkansas, Connecticut, Delaware, District of Columbia, Florida, Georgia, Illinois, Kentucky, Louisiana, Maine, Massachusetts, Mississippi, Missouri, New Jersey, New York, North Carolina, Oklahoma, Pennsylvania, Rhode Island, South Carolina, Texas, Virginia | 19316199 | Paulson (2017) |
| *Ischnura rufostigma* | FP(D) | P | BGD, HKG, LAO, MMR, NPL, THA, VNM | Assam, Bihar, Guangdong, Guizhou, Himachal Pradesh, Hunan, Madhya Pradesh, Manipur, Meghalaya, Nagaland, Sichuan, Uttarakhand, West Bengal, Yunnan | 4199316 | Mitra (2010), Sanmartin-Villlar et al. (2016) |
| *Ischnura saharensis* | FP(D) | P | DZA, LBY, MAR, MRT, NER, TUN | | 6717192 | Samraoui and Dijkstra (2010) |
| *Ischnura senegalensis* | FP(D) | P | AFG, AGO, ARE, BEN, BFA, BGD, BRN, BWA, CMR, COD, CPV, DJI, EGY, ESH, GAB, GHA, GMB, GNB, GNQ, HKG, IDN, IND, IRN, IRQ, ISR, JOR, JPN, KEN, KHM, LAO, LBR, LBY, LKA, LSO, MAC, MDG, MLI, MMR, MOZ, MRT, MUS, MWI, MYS, NAM, NER, NGA, OMN, PAK, PHL, PLW, PNG, PSE, REU, RWA, SAU, SDN, SEN, SGP, SLE, SOM, SSD, SWZ, SYC, TCD, TGO, THA, TKM, TLS, TWN, TZA, UGA, UZB, VNM, YEM, ZAF, ZMB, ZWE | Nei Mongol | 41440050 | Ozono et al. (2012), Sharma and Clausnitzer (2016) |
| *Ischnura* sp. | FM(H) | M | ASM | | 222 | Marinov et al. (2015) |

| Species | State | StateBin | Countries | Other divisions | Range area ($Km^2$) | Sources |
|---|---|---|---|---|---|---|
| *Ischnura taitensis* | FM(A) | M | PYF | | 3680 | Marinov et al. (2019) |
| *Ischnura ultima* | FP(D) | P | ARG, CHL | | 3537305 | von Ellenrieder and Garrison (2007), Heckman (2008) |
| *Ischnura verticalis* | FP(D) | P | CAN, USA | Arkansas, Colorado, Connecticut, Delaware, District of Columbia, Georgia, Illinois, Indiana, Iowa, Kansas, Kentucky, Maine, Massachusetts, Michigan, Minnesota, Missouri, Montana, Nebraska, New Brunswick, New Hampshire, New Jersey, New Mexico, New York, Newfoundland and Labrador, North Carolina, North Dakota, Nova Scotia, Ohio, Oklahoma, Ontario, Pennsylvania, Québec, Rhode Island, South Carolina, South Dakota, Tennessee, Texas, Vermont, Virginia, West Virginia, Wisconsin, Wyoming | 8679697 | Paulson (2009), Paulson (2012) |
| *Pacificagrion lachrymosa* | FM(H) | M | ASM | | 222 | Marinov et al. (2015) |
| *Pacificagrion* sp. | FM(H) | M | ASM | | 222 | Marinov et al. (2015) |

*Classification of female-colour states complemented by observation of field or museum specimens, as described in Willink et al. (2019)

**Table S3.** Geographic range areas for 23 species of *Ischnura* damselflies not included in this or previous studies (Robinson and Allgeyer, 1996; Mattern and van Gossum, 2008; Sánchez-Guillén et al., 2020). For each species, we report countries of known occurrence, with names abbreviated according to the International Organization for Standardization (ISO). We include additional information on geographic range such as names of islands, provinces or altitudinal range, where available. Geographic range information comes from literature listed under 'Sources'. Two species, *I. ordosi* and *I. rubella*, marked as "doubtful" in the World Odonata List (Schorr and Paulson, 2020) were excluded.

| Species | Countries | Other range information | Sources |
|---|---|---|---|
| *Ischnura acuticauda* | PNG | From middle to high elevations (1800-2900 m.a.s.l.) | Kalkman and Orr (2013) |
| *Ischnura albistigma* | WSM | Upolo Island and Tutuila Island | Fraser (1953) |
| *Ischnura ariel* | IDN | Papua, province, known from a few middle elevation sites (1740-1800 m.a.s.l.) | Kalkman and Orr (2013) |
| *Ischnura armeniaca* | IDN | Papua province, endemic to highlands (3200-3900 m.a.s.l.) | Kalkman and Orr (2013) |
| *Ischnura auricolor* | WSM | Upolo Island | Fraser (1927) |
| *Ischnura buxtoni* | WSM | Upolo Island | Fraser (1953) |
| *Ischnura chromostigma* | WSM | Tutuila Island | Fraser (1927) |
| *Ischnura filosa* | MDG | Madagascar Island | Fraser (1949a) |
| *Ischnura haemastigma* | WSM | Upolo Island | Fraser (1953) |
| *Ischnura inarmata* | IND | Kashmir | Calvert (1898) |
| *Ischnura isoetes* | IDN | Papua province, from high elevations (2500-3450 m.a.s.l.) | Kalkman and Orr (2013) |
| *Ischnura jeanyvesmeyeri* | PYF | Endemic to the island of Raivavae, Austral Islands | Englund and Polhemus (2010) |
| *Ischnura lorentzi* | IDN | Papua province, known from a single female | Kalkman and Orr (2013) |
| *Ischnura luta* | MNP | Endemic to island of Rota | Polhemus et al. (2000) |
| *Ischnura mahechai* | COL | Known from a highland lake (3600 m.a.s.l.) | Machado (2012) |
| *Ischnura oreada* | IDN | Papua province, endemic to highlands (3200-3700 m.a.s.l.) | Kalkman and Orr (2013) |
| *Ischnura rufovittata* | BOL | | Heckman (2008) |
| *Ischnura rurutana* | PYF | Endemic to the island of Rurutu, Austral Islands | Englund and Polhemus (2010) |
| *Ischnura sanguinostigma* | WSM | Upolo Island | Fraser (1953) |

| Species | Countries | Other range information | Sources |
|---|---|---|---|
| *Ischnura stueberi* | IDN | Papua province, known from several lowland localities | Kalkman and Orr (2013) |
| *Ischnura thelmae* | PYF | Endemic to the island of Rapa, Austral Islands | Englund and Polhemus (2010) |
| *Ischnura vinsoni* | MUS | Mauritius Island | Fraser (1949b) |
| *Ischnura xanthocyane* | IDN | Papua province, endemic to highlands (3000-3500 m.a.s.l.) | Kalkman and Orr (2013) |

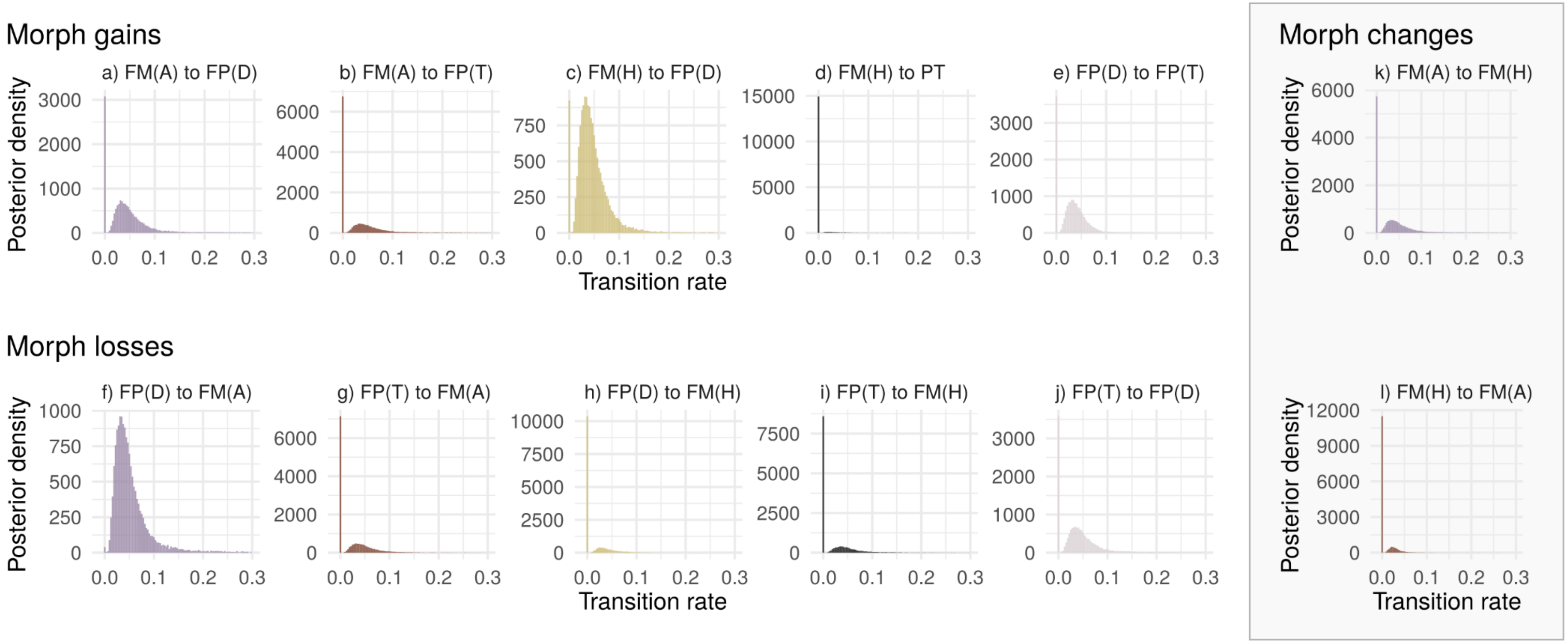


**Figure S1.** Posterior distribution of transition rate parameters in a multistate, reversible-jump (RJ) MCMC model of the evolution of female colour and female colour polymorphisms in *Ischnura* damselflies. The RJ model samples the parameter number and their values proportionally to their posterior probability. As a result, in each posterior sample a transition rate parameter may be equal to zero, or it may be included in a rate category with an estimated rate. Transition rate parameters may therefore have a bimodal posterior distribution, with a peak at zero and a peak at the mode of the posterior samples in which the estimated parameter is greater than zero. Transitions can occur between four character states: FM(A) = Monomorphic females with colour pattern similar to males (androchrome females), FM(H) = Monomorphic females with colour pattern markedly different from males (heterochrome females), FP(D) = Polymorphic females with two female morphs (one androchrome, one heterochrome), FP(T) = Polymorphic females with three female morphs (one androchrome, two heterochrome). **(a-e)** Transitions from FM(A) and FM(H) to either FP(D) or FP(T), and from FP(D) to FP(T) involve an evolutionary gain of novel female morphs. **(f-j)** Transitions away from FP(D) or FP(T) to FM(A) and FM(H), and from FP(T) to FP(D) occur if female morphs are lost. **(k-l)** Transitions in the grey box represent evolution of female or male colouration without the evolution of female polymorphisms.

**Supporting Materials and Methods**

*Expanded phylogeny reconstruction*

A recent study explored the origin and evolutionary dynamics of female colour polymorphism in *Ischnura* using a single MCC tree and stochastic character mapping (Sánchez-Guillén et al., 2020). The MCC tree incorporated nine species (*I. abyssinica, Ischnura sp. "a", I. aralensis, I. chingaza, I. cyane, I. forcipata, I. genei, I. indivisa, I. rubilio*) and two subspecies (*I. elegans ebneri* and *I. posita atezca*) that were additional to the subset used in this study. Sánchez-Guillén et al. (2020) inferred the most recent common ancestor (MRCA) of *Ischnura* as female-polymorphic (FP).

Sánchez-Guillén et al. (2020) conducted phylogenetic inference based on two mitochondrial and one nuclear marker, none of which were used in the present study due to poor taxonomic coverage. In order to investigate if our conflicting conclusions could be explained by the taxa missing in the present study, we used a Phylogenetic Assembly with Soft Taxonomic Inferences (PASTIS) approach (Thomas et al., 2013) in R v.3.6.1 (R Core Team 2020) to incorporate species lacking genetic data at the tree inference stage. As our MCC tree was inferred under a birth-death model, assuming each lineage is a species, we did not include the two subspecies sampled by Sánchez-Guillén et al. (2020) in these analyses. We used the PASTIS method to combine our MCC tree, inferred using StarBEAST2, sequence data (limited to one representative individual per species) and a set of taxonomic statements to guide the placement of the taxa lacking sequence data.

Each species, excluding *I. abyssinica* and *I. aralensis*, was treated as a separate clade, and missing taxa were assigned a number of sister species among which placement was random. We drew sister taxa assignments from the MCC tree in Sánchez-Guillén et al. (2020). In this guide tree, *I. abyssinica* and *I. aralensis* were inferred as sister species with 100% support. The phylogenetic placement of this two-species clade was uncertain in Sánchez-Guillén et al. (2020), so we restricted its placement in our extended tree to the least inclusive clade with support greater than 80%. Consequently, the sister taxa *I. abyssinica* and *I. aralensis* were randomly assigned within the clade in our MCC tree including *I. barberi, I. nursei, I. evansi, I. fluviatilis, I. fountaineae, I. saharensis, I. elegans, I. graellsii, I. rufostigma, I. ramburii, I. hetersticta, I. foylei* and *I. senegalensis* (see Fig. 2).

In the guide MCC tree (Sanchez-Guillén et al., 2020), three of the missing species (*I. cyane, I. indivisa* and *I. sp. "a"*) form a monophyletic clade with *I. caprelous,* and another species (*I. chingaza*) forms a monophyletic clade with *I. cruzi*. These clades are well supported at 100% support. Therefore, in our model we constrained the former three and latter one missing species to be sister taxa to *I. capreolus* and *I. cruzi* respectively. Equally, the missing species *I. rubilio* was inferred as sister to *I. aurora* at 100% in the guide MCC tree (Sanchez-Guillén et al., 2020). As Sánchez-Guillén et al. (2020) did not sample *I. pamelae* (inferred as sister to *I. aurora with* 100% support in the MCC tree from this study), *I. rubilio* was conservatively constrained to fall within the clade including *I. aurora* and *I. pamelae* in our PASTIS model. The placement of the missing species *I. forcipata* in a monophyletic clade with *I. asiatica, I. pumilio, I. ezoin* and *I. intermedia* was well-supported (99%) in the guide MCC tree, although the exact placement was uncertain (Sanchez-Guillén et al., 2020). Therefore, we placed this species randomly within this clade in our model.

Finally, we constrained the missing species *I. genei* to the clade in our MCC tree including *I. nursei, I. evansi, I. fluviatilis, I. fountaineae, I. saharensis, I. elegans, I. graellsii, I. rufostigma, I. ramburii, I. hetersticta, I. foylei* and *I. senegalensis.* In Sánchez-Guillén et al. (2020), *I. genei* falls within a less inclusive clade with relatively high support (82%). Yet, this less inclusive clade does not appear in the MCC tree from the present study. We thus assigned a random phylogenetic placement of *I. genei* to a more inclusive clade that is shared by Sánchez-Guillén et al. (2020) and the present study. However, this clade is not strongly supported (< 80%) in Sánchez-Guillén et al. (2020), and has moderate support in this study (79%).

Bayesian assessment of the placement of missing taxa was then conducted using MrBayes v.3.2.7 (Ronquist et al., 2012). Most of the default settings outlined in the PASTIS template were used in the phylogenetic model and MCMC algorithm implemented in MrBayes. We placed a birth-death prior on the species tree and fixed extinction rates to zero. We used an independent gamma rate (IGR) relaxed clock model and the expected variability in branch lengths was drawn from an exponential prior with a mean of 10. We also placed an exponential prior with a mean of 1 on the speciation rate. We specified a molecular evolution model with 5 partitions, one for each locus. Each partition had its own generalized time reversible (GTR) substitution model under default priors. Site-rate heterogeneity was modelled as gamma distributed with four rate categories for all loci, except for D7, in which the proportion of invariant sites was estimated. A unique rate multiplier was set for each partition, except for the two mitochondrial markers (16S and COI), which were linked. We ran a single Markov Chain Monte Carlo (MCMC) algorithm for 50 million iterations with a sampling frequency of 10000 and disposed of the first 10 million samples as burnin. Stationarity and convergence checks of MCMC runs were performed as per the main text (see Materials and Methods).

Out of the nine additional species included in this analysis, two (*I. rubilio* and *I. forcipata*) were classified as female-monomorphic with heterochrome females (FM(H)) according to Sánchez-Guillén et al. (2020). Six species (*I. aralensis*, *I. abyssinica*, *I. indivisa*, *I. sp. a*, *I. cyane* and *I. chingaza*) were classified as female-polymorphic with dimorphic females (FP(D)) (Sánchez-Guillén et al. 2020). However, we note that for *I. cyane* and *I. chingaza* this classification was based on unpublished observations, and a taxonomic study cited by Sánchez-Guillén et al. (2020) indicates that colour polymorphisms have not been found so far in these species (Realpe, 2010). This also is the case for *I. cruzi*, which we classified as FM(H), but was classified as FP(D) in Sánchez-Guillén et al. (2020), based on unpublished observations. For this analysis we assumed *I. cruzi* to be FP(D) as in Sánchez-Guillén et al. (2020). *I. genei,* the last missing taxon, was classified as female-polymorphic with trimorphic females (FP(T)) (Sánchez-Guillén et al., 2020).

There are four other conflicts in female-colour classification between the present study and Sánchez-Guillén et al. (2020). We classified *I. aurora* as FP(D), as androchrome females are described and photographed, albeit reportedly rare, in Ozono et al. (2012). In contrast, this species was classified as FM(H) in Sánchez-Guillén et al. (2020). *I. evansi* was classified as FP(T) by Sánchez-Guillén et al. (2020) and FP(D) in the present study, as we could not ascertain if the three phenotypic forms in Marzoq (2005) represented one androchrome and two heterochrome morphs or one androchrome and one heterochrome morph sampled in two developmental colour phases. Sánchez-Guillén et al. (2020) classified *I. posita* as FM(H). Here, we classified it as FM(A), according to Paulson (2012), who described females as similar in colour pattern as males, but with postocular spots and thorax stripes pale to blue rather than light green, and developing more pruinosity with age than males do. Finally, *I. heterosticta* was classified as FM(H) in Sánchez-Guillén et al. (2020), citing Huang et al. (2012). However, Huang et al. (2012) actually describe two distinct female morphs, an andro-gynochrome morph that starts out with a male-like colour pattern and changes to a green and then grey colour with age, and a gynochrome (same as heterochrome) morph that does not undergo this ontogenetic colour change.

Of these four conflicts, two (*I. aurora* and *I. heterosticta*) affect whether a species is classified as female polymorphic (FP) or female monomorphic (FM), whereas the other two only relate to the extent of sexual dimorphism in a FM taxon (*I. posita*), or the number of female morphs in a female-polymorphic species (*I. evansi*). As the analysis in Sánchez-Guillén et al. (2020) only considered whether females were monomorphic (FM) or polymorphic (FP), and we re-classified *I. cruzi* as FP(D), only 2 out of 50 species have a directly conflicting female-colour classification between this supplementary analysis and the one in Sánchez-Guillén et al. (2020).

We then used the Multistate method in BayesTraits v.3.0 (Pagel et al., 2004) to infer female-colour character states at ancestral nodes, given the extended *Ischnura* phylogeny and the female-colour character states of extant species. We ran the Multistate model using a sample of 1000 trees from one independent run of the phylogenetic assembly analyses and kept all phylogenetic model and MCMC algorithm parameters the same as per the analyses from the main text (see Materials and Methods). Ancestral character states were mapped onto the maximum clade credibility (MCC) tree, drawn from the MrBayes output.

### Results

The results from this analysis were qualitatively similar to the main ancestral state reconstruction of this study, and the MRCA of *Ischnura* was again inferred as being FM(H). However, the female-colour state was inferred with a slightly lower mean posterior probability (PP) than in the main analyses of 66% (Fig. S2). Equally, the probability that the ancestor was FP(D) increased to 33%. We note that 7 out of the 9 previously missing taxa included in these analyses are either FP(D) or FP(T), likely driving the increase in probability of a FP ancestor in *Ischnura.* However, FM species may be just as numerous as FP species across the genus (Willink et al. 2019), and they remain under-sampled as they are more likely to occupy confined geographic ranges such as remote islands (see Discussion).

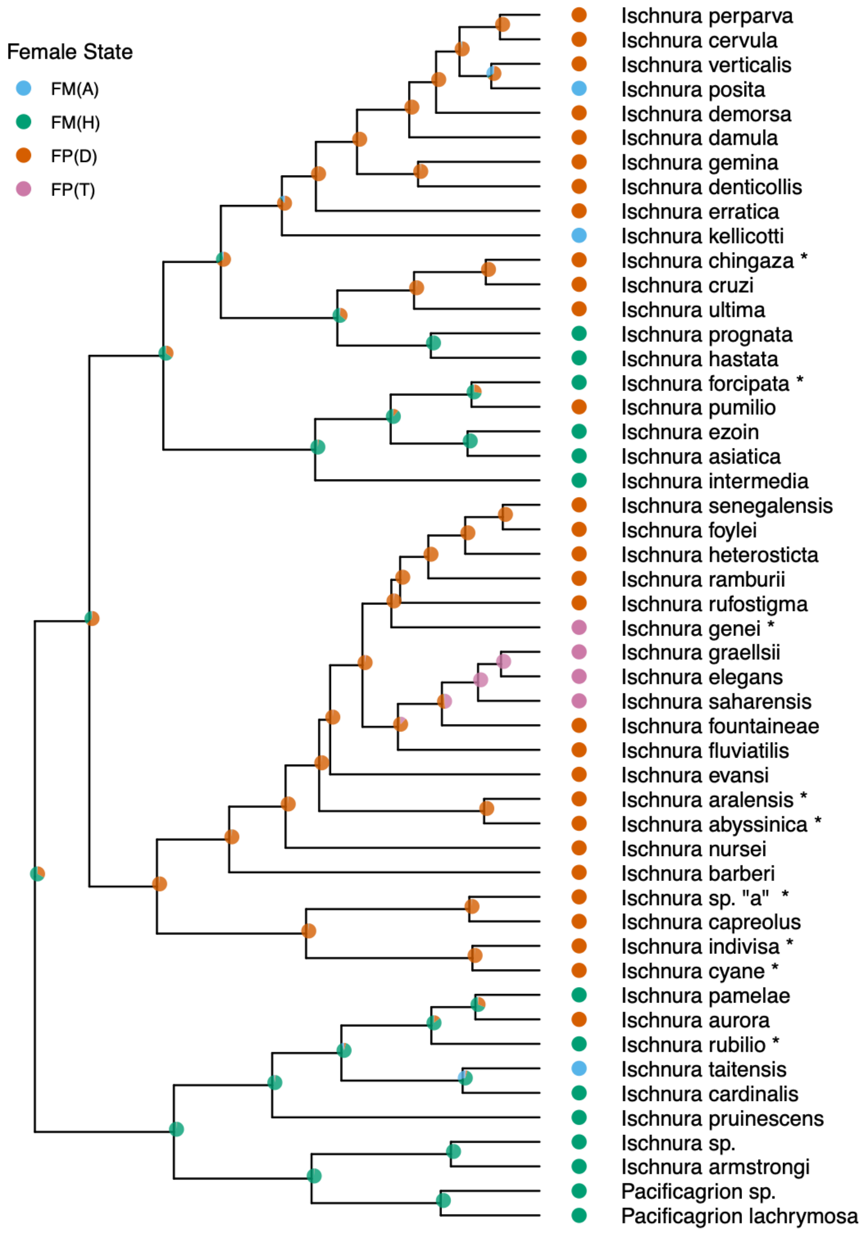


**Figure S2.** Ancestral state reconstruction plotted on the extended *Ischnura* MCC tree. Nine additional species (labelled with asterisks) were added using soft taxonomic inferences. Ancestral state reconstruction was conducted on a posterior sample of 1000 trees. For each tree, the positions of missing taxa were constrained to specific clades, based on previous phylogenetic inference, and randomized within those clades (see Supporting Methods). We plot these results on a single MCC tree for simplicity. Extant morph states are represented by circles at tips and reconstructed ancestral states are represented by piecharts at nodes (female-heterochromatic only, green; female-andromorphic only, blue; female-dimorphic, red; female-trimorphic, pink). Piecharts show the mean posterior probability for each morph state and are plotted on the consensus tree.
